# Modelling Eurasian lynx populations in Western Europe: What prospects for the next 50 years?

**DOI:** 10.1101/2021.10.22.465393

**Authors:** Bauduin Sarah, Germain Estelle, Zimmermann Fridolin, Idelberger Sylvia, Herdtfelder Micha, Heurich Marco, Kramer-Schadt Stephanie, Duchamp Christophe, Drouet-Hoguet Nolwenn, Morand Alain, Blanc Laetitia, Charbonnel Anaïs, Gimenez Olivier

**Affiliations:** Office français de la biodiversité, 147 avenue de Lodève, 34990 Juvignac, France; Centre de Recherche et d’Observation sur les Carnivores, 57590 Lucy, France; Carnivore Ecology and Wildlife Management. KORA, Thunstrasse 31, CH-3074 Muri bei Bern, Switzerland; Department of Ecology and Evolution, University of Lausanne, Lausanne, Switzerland; Stiftung Natur und Umwelt Rheinland-Pfalz, Diether-von-Isenburg-Str. 7, 55116 Mainz, Germany; Forest Research Institute of Baden-Wuerttemberg, Wonnhalde 4, 79100 Freiburg, Germany; Chair of Wildlife Ecology and Management, Faculty of Environment and Natural Resources, University of Freiburg, Tennenbacher Straße 4, 79106 Freiburg, Germany; Department of National Park Monitoring and Animal Management, Bavarian Forest National Park, Freyunger Straße 2, 94481 Grafenau, Germany; Faculty of Forestry and Wildlife Management, Campus Evenstad, University of Inland Norway, 2480 Koppang, Norway; Department of Ecological Dynamics, Leibniz Institute for Zoo and Wildlife Research, D-10315 Berlin, Germany; Institute of Ecology, Technische Universität Berlin, Germany; Office français de la biodiversité, Micropolis - La Bérardie F-05000 Gap, France; Office français de la biodiversité, 5 Allée de Bethléem, F-38610 Gières, France; Centre d’études et d’expertise sur les Risques, l’Environnement, la Mobilité et l’Aménagement, Direction territoriale Est, Ile du Saulcy, 57045 Metz, France; CEFE, Univ Montpellier, CNRS, EPHE, IRD, Montpellier, France

**Keywords:** *Lynx lynx*, population persistence, spatially-explicit individual-based model

## Abstract

Persistence of populations may be uncertain for large carnivore species, especially for those established in human-dominated landscapes. Here, we studied the Eurasian lynx in Western Europe established in the Upper Rhine meta-population (i.e., Jura, Vosges-Palatinian and Black Forest populations) and the Alpine population. These populations are currently considered endangered or critically endangered due to high anthropogenic mortality, small population size and low genetic diversity, and isolation. We assessed lynx persistence over a 50-year time horizon by implementing a spatially-explicit individual-based model, while accounting for road mortality and habitat selection. Forecasts showed a steady growth rapidly reaching a more stable phase for the Alpine and Jura populations, and a more heterogeneous positive growth with less precision for the Vosges-Palatinian and Black Forest populations. Exchanges of individuals between populations were limited, the Jura population playing the role of a crossroad. Finally, the persistence of lynx in Western Europe seems likely on a large scale over the next 50 years. Indeed, simulations showed high female occupancy as well as average lynx density over the core areas of the four studied populations. Nevertheless, these results should be interpreted with the model limitations in mind, concerning the absence of movement barriers and inbreeding depression.

## Introduction

The Eurasian lynx (*Lynx lynx*) was eradicated in most of Europe between the 17^th^ and 20^th^ centuries. The main reasons for its disappearance were habitat degradation, human persecution and a decrease in prey availability (Breitenmoser et al., 2000). The species has recently recolonized parts of its historical range in Central and Western Europe thanks to different reintroduction programs which started in the 1970s. Nowadays, there are eleven identified lynx populations in Europe (von Arx, 2020), and the species benefits from a conservation status across its whole range area. The species is considered as “least concerned” at the European level of the IUCN Red list. However, its status greatly differs from one population to another, notably because of their difference in population sizes, even though these populations share similar threats, mostly habitat destruction and fragmentation, illegal killings and collisions with vehicles (von Arx, 2020). Long-term persistence remains notably uncertain for the Alpine population (France and Switzerland) and for the Upper Rhine meta-population, which encompasses the Jura population (France and Switzerland), the Vosges-Palatinian population (France and Germany) and few individuals located in the Black Forest and its surroundings (e.g., the Swabian Alb, Germany) (Drouet-Hoguet et al., 2021; Germain & Schwoerer, 2021; Herdtfelder, Schraml, & Suchant, 2021; Idelberger et al., 2021; Molinari-Jobin et al., 2021). These populations are currently defined as endangered (Jura and Alpine) or critically endangered (Vosges-Palatinian) (von Arx, 2020). Indeed, the populations forming the Upper Rhine meta-population remain currently small and isolated. Only a few exchanges between them and with the Alpine population are documented, most likely because of habitat fragmentation and little functional connectivity (Zimmermann & Breitenmoser, 2007; Morand, 2016). Low female dispersal rates and movements most likely slow population expansion due to their conservative dispersal behavior compared to male lynx (Port et al., 2021). This situation, added to the common origin from the Carpathian population (Breitenmoser, Breitenmoser-Würsten, & Capt, 1998; Vandel et al., 2006) of the few reintroduced individuals which did reproduce, resulting in a bottleneck and genetic drift, may lead to high inbreeding within populations (Breitenmoser-Würsten & Obexer-Ruff, 2003; Bull et al., 2016; Premier et al., 2020). A national action plan has been defined in France with several objectives to restore the lynx to a favorable conservation status in France (Gatti, 2022).

In this context, wildlife conservationists, scientists and policy-makers face significant challenges for lynx conservation when individuals inhabit human-dominated landscapes. Several studies have used individual-based models to inform decision-making processes for population conservation (IBMs; Railsback & Grimm, 2012). These bottom-up models flexibly integrate species demography (dispersal, territory establishment, reproduction, mortality) and how animals interact with their inhomogeneous environment (e.g., habitat selection, collisions, illegal killings) and other individuals, while accounting for individual characteristics (e.g., sex, age, dispersal status). Population-level results emerge from the individual-level simulations (Railsback & Grimm, 2012). These models are used for the management and conservation of large and small carnivores (e.g., Kramer-Schadt et al., 2004, 2011; Kramer-Schadt, Revilla, & Wiegand, 2005; Marucco & McIntire, 2010; Hradsky et al., 2019). Recent applications of IBMs for the lynx species allowed estimating illegal killing (Heurich et al., 2018), linking movements and genetic diversity (Premier et al., 2020) and evaluating reintroduction scenarios (Ovenden et al., 2019; Philips, 2020).

Lynx are large carnivores with extensive spatial requirements (Breitenmoser-Würsten et al., 2007). They are territorial and live in vast connected forested areas (Vandel & Stahl, 2005; Zimmermann & Breitenmoser, 2007; Ripari et al., 2022), which sustain important prey populations (Basille et al., 2009). Natural barriers (e.g., broad waterbodies) as well as anthropogenic barriers (e.g., urban areas, road network) limit lynx movements (Kramer-Schadt et al., 2004). Roads are complex infrastructures acting as semi-permeable movement barriers (Klar, Hermann, & Kramer-Schadt, 2006; Zimmermann & Breitenmoser, 2007; Bastianelli et al., 2024) in addition to being an important source of mortality (Schmidt-Posthaus et al., 2002; Ceia-Hasse et al., 2017; Bencin et al., 2019). Therefore, a major modelling challenge when simulating lynx populations is to account for movements over large areas and consequently, for the impact of landscape characteristics on different demographic processes such as dispersal, territory establishment and survival rates (Premier et al., 2025). Simulation rules in IBMs are defined at the individual level and result in different behavior of each simulated individual due to their characteristics. Therefore, these models appear to be a great tool to integrate the complex interactions between landscape characteristics and both lynx demographic and spatial processes, to simulate population changes on a large scale.

Our study aimed to assess the long-term persistence of the Eurasian lynx in the large range area delimited by the Upper Rhine meta-population and the Alpine population, using the available population status in 2018. We build a spatially-explicit IBM that integrates up-to-date information on lynx ecology while accounting for habitat preferences and the risk of collisions with vehicles. We implement our model in the R programming language and share it publicly with open-source code allowing full reproducibility and its adoption by ecologists and conservationists. We use this model to forecast the fate of lynx populations over a 50-year time horizon. Finally, we provide several population metrics that help in diagnosing the long-term species persistence in its most western range area.

## Material and Methods

### Study area and populations

We conducted the study on the Eurasian lynx populations located in France, Germany and Switzerland (Fig. 1). The three main populations inhabiting the vast mountainous and forested areas of these countries are the Vosges-Palatinian (France-Germany), the Jura (France-Switzerland) and the Alpine (France-Switzerland) populations (von Arx, 2020). Some individuals were also observed in the Black Forest and its surroundings (e.g., Swabian Alb) in Germany (Fig. 1), but were not considered as a lynx population *per se* (von Arx, 2020). Therefore, we should refer to “mountain ranges” when speaking of the Vosges-Palatinate, Jura, Alps and Black Forest with its surroundings, but for simplicity and clarity, we use “populations” throughout our paper.

**Figure 1:**
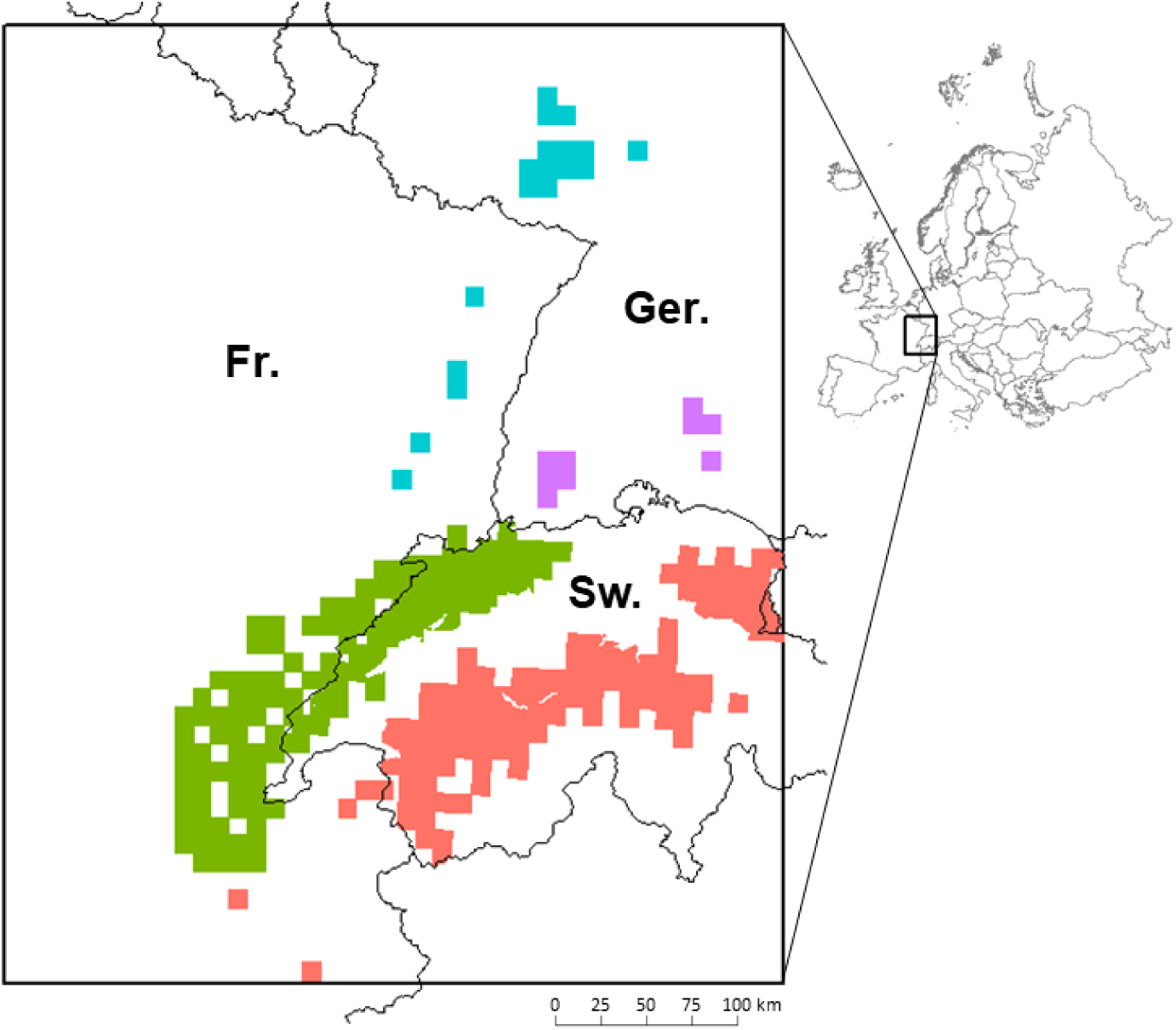
Eurasian lynx presence as available in 2017-2018 in France (Fr.), Germany (Ger.) and Switzerland (Sw.) over the study area (black rectangle). Data for France cover the period from 01/04/2013 to 31/03/2017 (OFB Réseau Loup-Lynx), for Germany from 01/05/2017 to 30/04/2018 (Bundesamt für Naturschutz) and for Switzerland from 01/01/2015 to 31/12/2017 (KORA). We used a standardized 10×10 km grid from the European Environment Agency for France and Germany, and a grid derived from the 1:25,000 map for Switzerland). The four colors are for the four different populations: the Vosges-Palatinian population (blue; 1,800 km^2^) with cells in France (Vosges Mountains) and Germany (Palatinate Forest), the Black Forest population (purple; 900 km^2^) in Germany, and the Jura (green; 12,057 km^2^) and the Alpine (red; 11,190 km^2^) populations with cells in France and Switzerland. Top right corner inset: Europe with the location of the study area.

The Jura and Alpine populations originated from individuals reintroduced in Switzerland in the 1970s (Breitenmoser, Breitenmoser-Würsten, & Capt, 1998; Vandel & Stahl, 2005), followed by a natural recolonization of the territories. After the complete decline of the Vosges-Palatinian population in the 18^th^ century, a reintroduction of 21 lynx occurred in the southern part of the Vosges Mountains (France) between 1983 and 1993 (Vandel et al., 2006). Only 10 individuals contributed to the local establishment of the species (Vandel et al., 2006), without conclusive stabilization (Charbonnel & Germain, 2020). Since 2016, the lynx has been back in the Palatinate Forest (Germany), thanks to a new reintroduction program finalized in 2020, with a few individuals that already moved to the Vosges Mountains (Scheid, Germain, & Schwoerer, 2021; Schwoerer, 2021). Finally, only a few male individuals have been observed in the Black Forest area since 2013, most of them coming from the Swiss Jura Mountains (Stiftung KORA, 2017; Ministerium für Ländlichen Raum und Verbraucherschutz, 2019; Wölfl et al., 2021).

### Lynx population model

Based on previous works by Kramer-Schadt et al. (2004, 2005, 2011), we built a spatially-explicit individual-based model (SE-IBM) to simulate lynx populations dynamics and dispersal, while accounting for the risk of lynx-vehicle collisions and lynx habitat preferences. An SE-IBM is an IBM where individual responses to behavioral rules are constrained by environmental characteristics. Our lynx SE-IBM is made of four components (Fig. 2) for which the complete description is available in Appendix A. The first component represents the impact of the road network on lynx survival. Hereby, the “collision layer” (Fig. A.1) represents lynx-vehicle lethal collision probabilities and is based on a collision model. The second component represents the impact of land cover on lynx space use with a habitat model yielding a “habitat layer” (Fig. A.2) and showing the distribution of the different lynx habitat types. These two spatial layers influence the behavioral rules followed by the simulated lynx individuals. The third component represents the initial lynx populations made of lynx individual’s location (Fig. A.3) and characteristics (e.g., age, sex). The fourth component details all SE-IBM rules including lynx demography, dispersal movement and territory establishment. The four components were combined to create an SE-IBM and simulate lynx populations. The initial lynx populations are the starting individuals created at the beginning of the simulation. Simulated individuals are either resident (i.e., established with a territory) or dispersers (Fig. A.4). Dispersing lynx move through the landscape, driven by a preference towards certain habitat types defined by the “habitat layer”, a constant direction and their memory (Fig. A.5). Dispersers can die with a fixed annual mortality probability or from vehicle collisions (additive spatial mortality), defined using the “collision layer”. When arriving at a new location, dispersers look for a territory (Fig. A.6). Females need a large area of good quality habitat to establish, whereas male establishment is driven by the presence of available resident females. A female resident may reproduce if there is a resident male available nearby.

**Figure 2:**
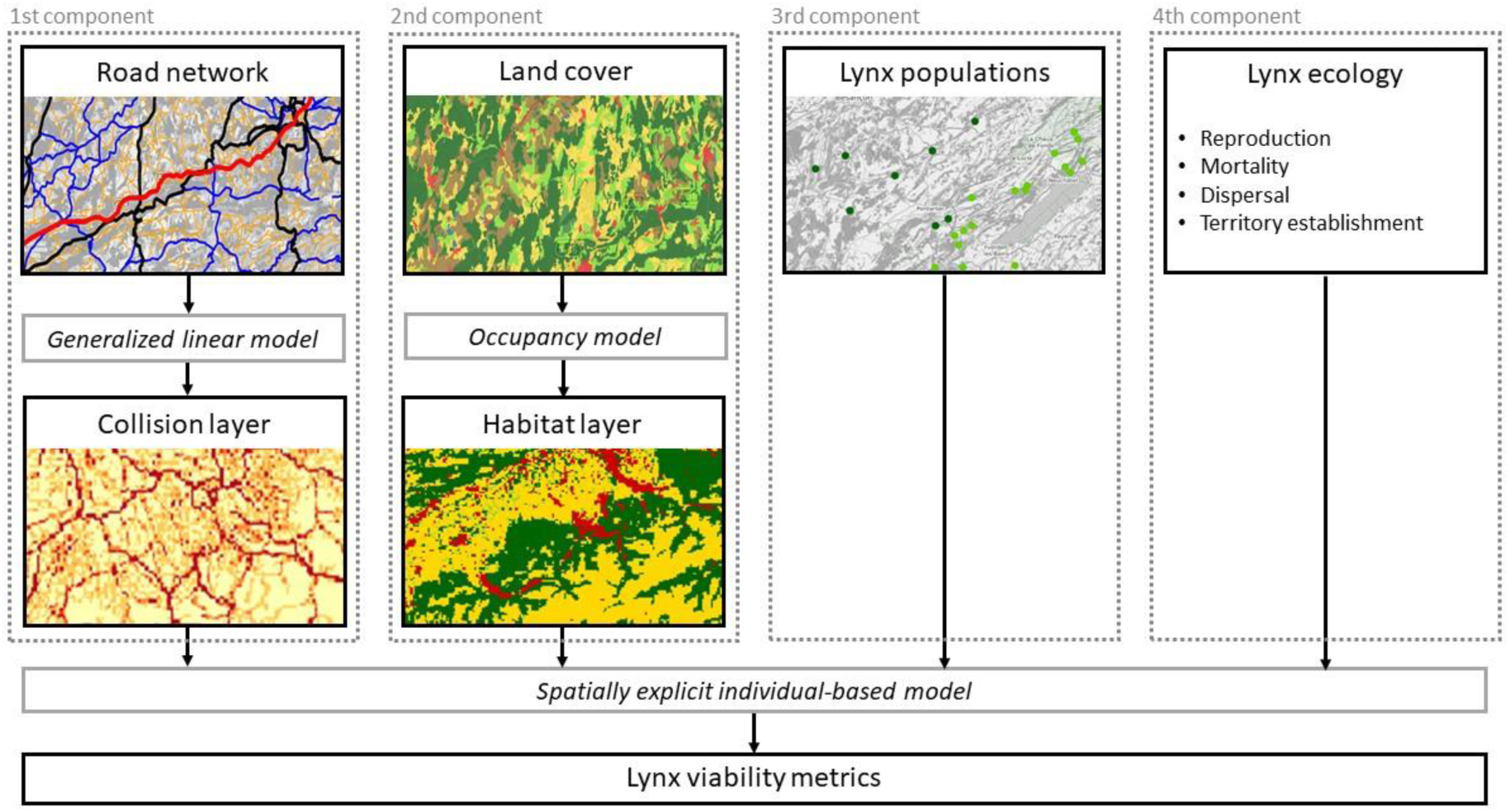
The four components of the lynx SE-IBM. The first component represents the impact of the road network on lynx survival through vehicle collision. A generalized linear model predicts collision probabilities (“Collision layer”, Fig. A.1). The second component represents the impact of land cover on the lynx populations. A site-occupancy model predicts lynx habitats influencing lynx movement (“Habitat layer”, Fig. A.2). The third component represents lynx populations with individuals’ locations and characteristics (Fig. A.3). The fourth component encompasses all ecological rules and SE-IBM parameters. All four components are included in the SE-IBM to simulate lynx populations and assess their persistence with different metrics.

Residents may also die with a fixed mortality or by vehicle collision. This sequence represents one year and is repeated with the updated individuals (e.g., newborns added, dead individuals removed, individuals at different locations, new residents) as long as desired. A complete description of the SE-IBM structure and processes following the Overview, Design concepts, and Details (ODD) protocol (Grimm et al., 2006, 2010) is provided in Appendix B. The full list of model parameters with their values is also provided in Appendix B (Table B.1). We calibrated parameters to fit the model to our studied populations better. We also performed a sensitivity analysis to evaluate the changes in the main results following the changes in parameter values. Full details on model calibration, sensitivity analysis and validation are available in Appendix C.

### Population persistence

We ran 100 replicates of the lynx SE-IBM to forecast the populations over 50 years. We used a different initial population (Appendix A) at each replicate to avoid bias due to the initial locations of simulated individuals. Initial populations were created based on real monitoring data (Appendix A). We defined a burn-in phase of 2 years after the start of the simulation (i.e., only simulation outputs from the 3^rd^ year of simulation were included in the conclusions of the analyses) to let the population settle down as all individuals from the initial population were defined as dispersers. We did not simulate any mortality during the first year of simulation to allow all individuals to find a territory, if they can, without dying. To evaluate lynx persistence, we analyzed a) population growth rates, b) number of movements between populations looking at establishments outside of the lynx native population, c) female territory occupancy, and d) lynx density.

a. We extracted the number of individuals in each population, for every year and each replicate. Then, we calculated the growth rate for each replicate and each population over the simulated years as the number of individuals at time *t* divided by the number of individuals at time *t*-1. Finally, we calculated the mean growth rate per year and population, and the 95% confidence interval of the mean over the 100 replicates.
b. We extracted the number of times individuals established their territory in a population area not corresponding to the one they were born in (“Population layer”, Appendix A) for every year and for each replicate. We did not account for individuals which moved to another population area during their dispersal but finally came back to their native population area to establish, nor those which died in another population area while dispersing. We calculated the number of lynx establishing outside of their natal population per year for each replicate and direction (e.g., from the Alpine to the Jura population and vice versa, from the Jura to the Vosges-Palatinian population and vice versa). Finally, we calculated the mean and 95% confidence interval of the mean across all replicates.
c. We extracted female territory locations at the last year of simulation for each replicate. We focused on female territories as male territories are based on those of the females. We assigned a “1” to each cell of the gridded study area (1 km^2^) that was occupied by a female territory and a “0” otherwise. Finally, we estimated territory occupancy by calculating the mean value per cell from these rescaled maps over the 100 replicates. We also visually validated our model predictions using GPS and VHF tracks from several collared female residents. To this end, we overlaid telemetry locations on the resulting territory occupancy map to see if they mostly fall in cells often selected by simulated females, to establish their territory (details in Appendix D).
d. We extracted the locations of all adult residents in the last year of simulation for each replicate. As density measure, we computed the mean number of lynx on each 1 km^2^ cell over all replicates. Finally, we summed these means at the 100 km^2^ resolution cells to obtain lynx density values at this scale.

## Results

### Population growth rates

Simulations predicted similar growth rate patterns for all lynx populations, with growth rates above one (i.e., growing phase) slowly decreasing towards reaching one (i.e., stabilization phase at saturation) along the 50 years of simulation (Fig. 3). The Alpine and Jura populations had similar patterns with their maximum growth rates equal to 1.09 (sd = 0.04) and 1.05 (sd = 0.06) at the 10^th^ and 13^th^ year of simulation, respectively. The Jura population quickly reached a stabilization phase around the 20^th^ year and fluctuated a little over the final period of the simulation. The Alpine population growth rate similarly decreased rapidly, reaching a low value, but still significantly over 1. The Vosges-Palatinian and Black Forest populations had more fluctuating growth rates over the simulation. It reached a maximum of 1.18 (sd = 0.51) and 1.45 (sd = 0.93) at the 23^rd^ and 14^th^ year of simulation, respectively. Confidence intervals around mean growth rates were much larger than those of the Alpine and Jura populations. The Vosges-Palatinian and Black Forest populations also had their growth rates decreasing after reaching their maximum, until reaching values close to 1 but only towards the end of the simulation period.

**Figure 3:**
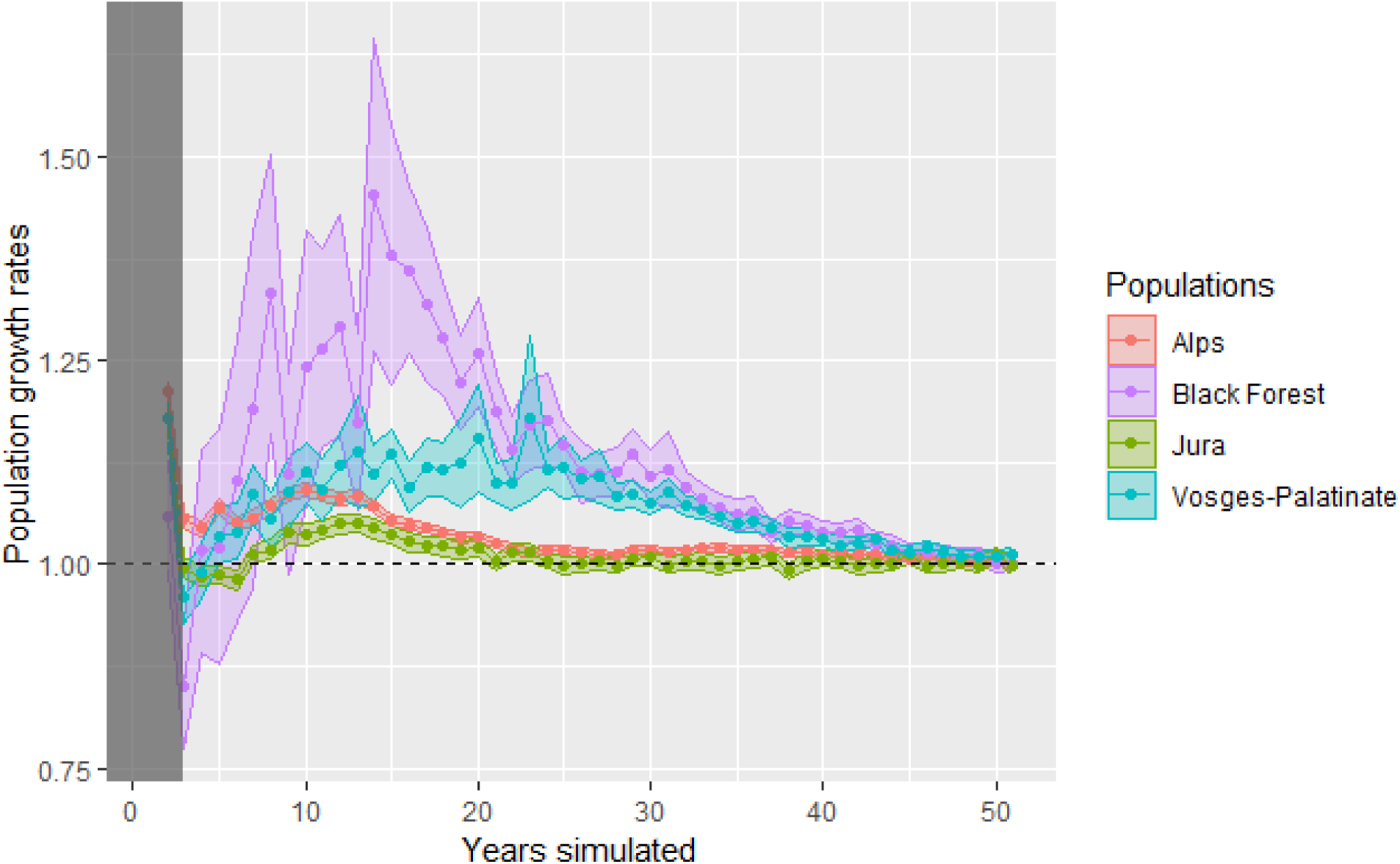
Annual rates of increase over the simulated years for each population. The grey area represents the 2-year burn-in phase at the beginning of the simulation. Points are means over 100 replicates and colored envelopes are 95% confidence intervals from 100 replicates. The dashed line represents a stabilization of the population (growth rate equals to 1).

### Lynx establishments outside their native populations

The number of lynx establishments in a population area different from their native one showed that movements between populations were rare for the four populations. Indeed, the highest values were 2.5 lynx per year on average, concerning individuals born in the Alps and successfully established in the Jura when reaching 25 years of simulation (Fig. 4.a). The Jura population seemed to be at the center of lynx movements among populations in Western Europe (Fig. 4). However, the number of individuals born in Jura and establishing elsewhere decreased around the 20^th^ year of simulation, matching when the population started to stabilize (Fig. 3 and 4.b). There was almost no exchange of individuals between the Alpine and Vosges-Palatinian populations, in any direction (Fig. 4.a and 4.c); as these are the most distant populations, the other ones must serve as stepping stones. Surprisingly, the Black Forest showed an exchange of individuals mainly with the Vosges-Palatinian population, even though the Black Forest seemed evenly distant to all three other populations (Fig. 4).

**Figure 4:**
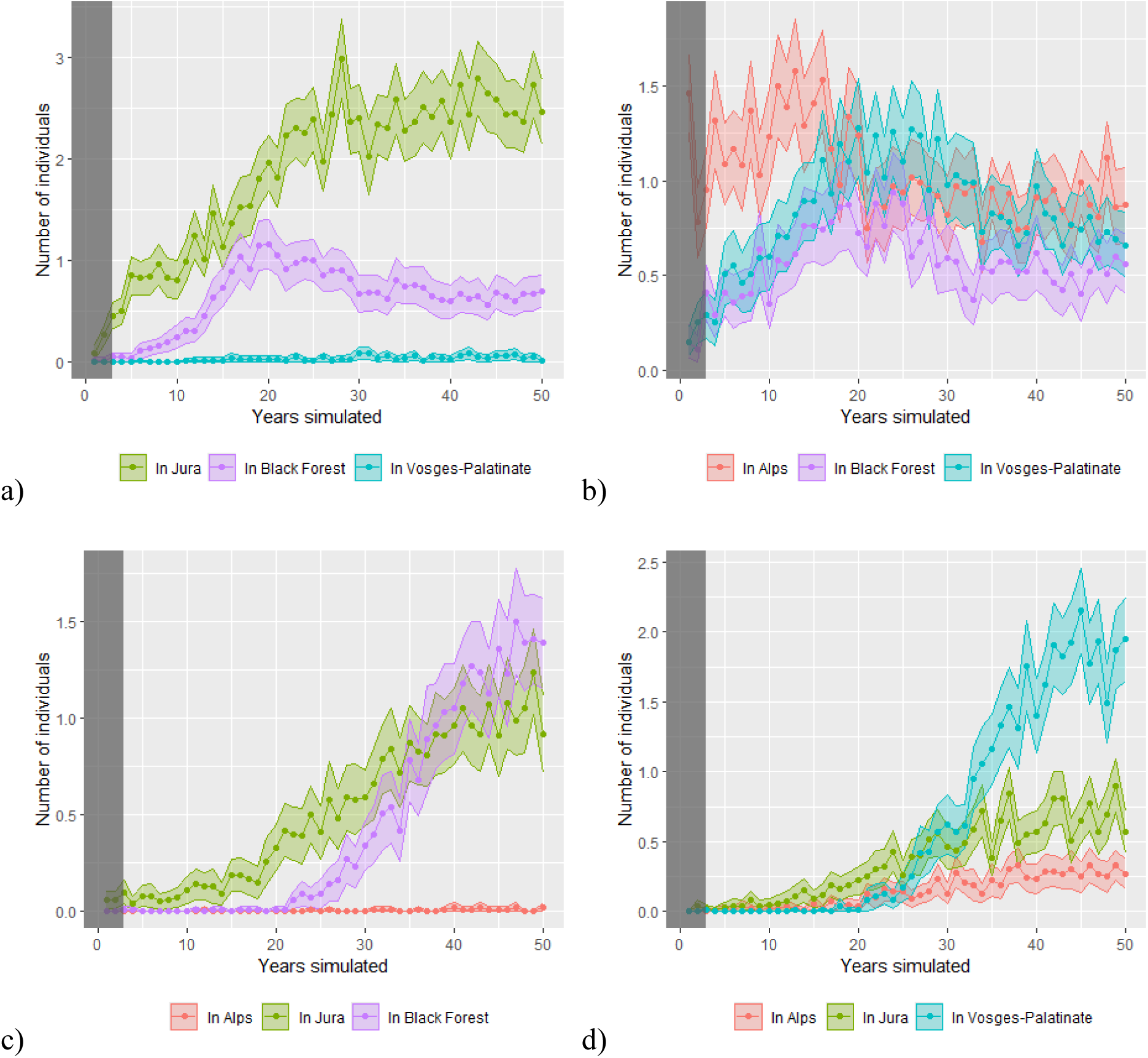
Number of lynx establishments outside their native populations per year when born in the a) Alpine, b) Jura, c) Vosges-Palatinian or d) Black Forest population. Colors represent different directions from the native populations (each figure) to those where individuals established (different colored lines). The grey area represents the 2-year burn-in phase. Points are mean over 100 replicates and colored envelopes are 95% confidence intervals around the mean from 100 replicates.

### Territory occupancy

At the end of the simulation period, female territories occupied most of the study area (Fig. 5). In the core areas of the four populations, female occupancy was high, reaching values above 0.85. The overlaid VHF and GPS paths of collared resident females (Fig. 5 and Appendix D) matched well the limits of the most frequently occupied cells. Most patches far from these core areas were never occupied over the 100 replicate simulations (Fig. 5). Nevertheless, there were distant patches, especially the bigger ones, occupied by female territories with low probabilities (Fig. 5).

**Figure 5:**
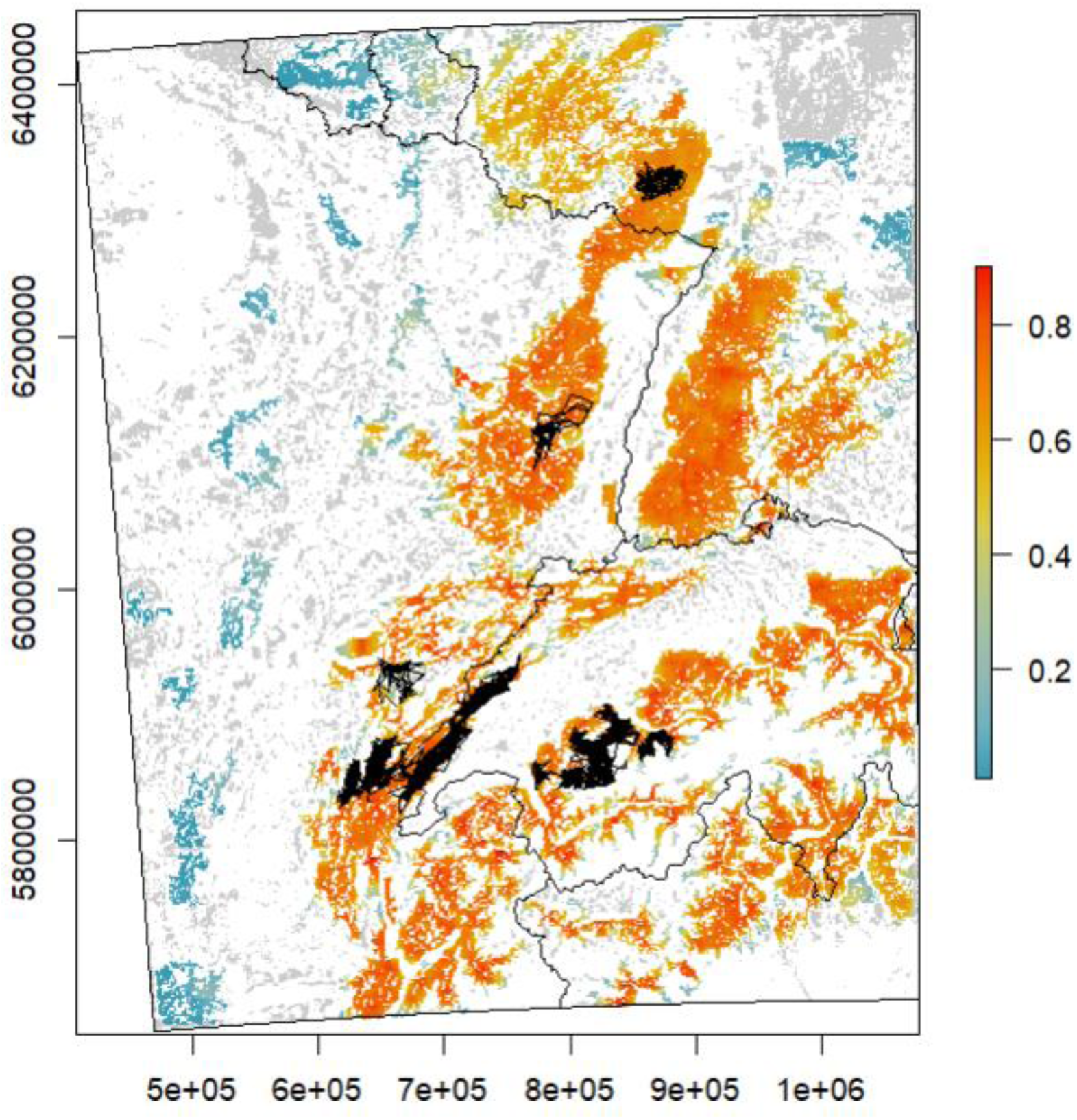
Occupancy by female territories over the study area in the last year of simulation. Values, ranging from zero (excluded) to one represented by the color scale blue-yellow-red, are mean occupancy probability per cell of 1 km^2^ over 100 replicates (e.g., a cell equal to 1 means the cell was included in a female territory, at the 50^th^ year of simulation, in all 100 simulation replicates). Grey cells are “breeding habitats” never included in a female territory across all replicates (equal to zero). White cells are cells of a habitat type different than “breeding habitat” and therefore could never be included in a female territory. GPS and VHF recorded paths for female residents are overlaid as thin black lines. A zoom of the different areas with telemetry data is presented in Appendix D. Axis (x and y) are in meters from the WGS 84 / Pseudo-Mercator coordinate system.

### Lynx density

Adult resident density ranged from 0 to 1.69 lynx per 100 km^2^ over the study area, in the last year of simulation (Fig. 6). The highest density value was in the Alpine population and this population also had the highest mean density over the four populations (0.55, sd = 0.4). The Jura population had the lowest density values (max = 0.97, mean = 0.26, sd = .024). The Vosges-Palatinian and Black Forest had intermediate values (max = 1.13 and 1.16, mean = 0.33 and 0.47, sd = 0.29 and 0.34, respectively).

**Figure 6:**
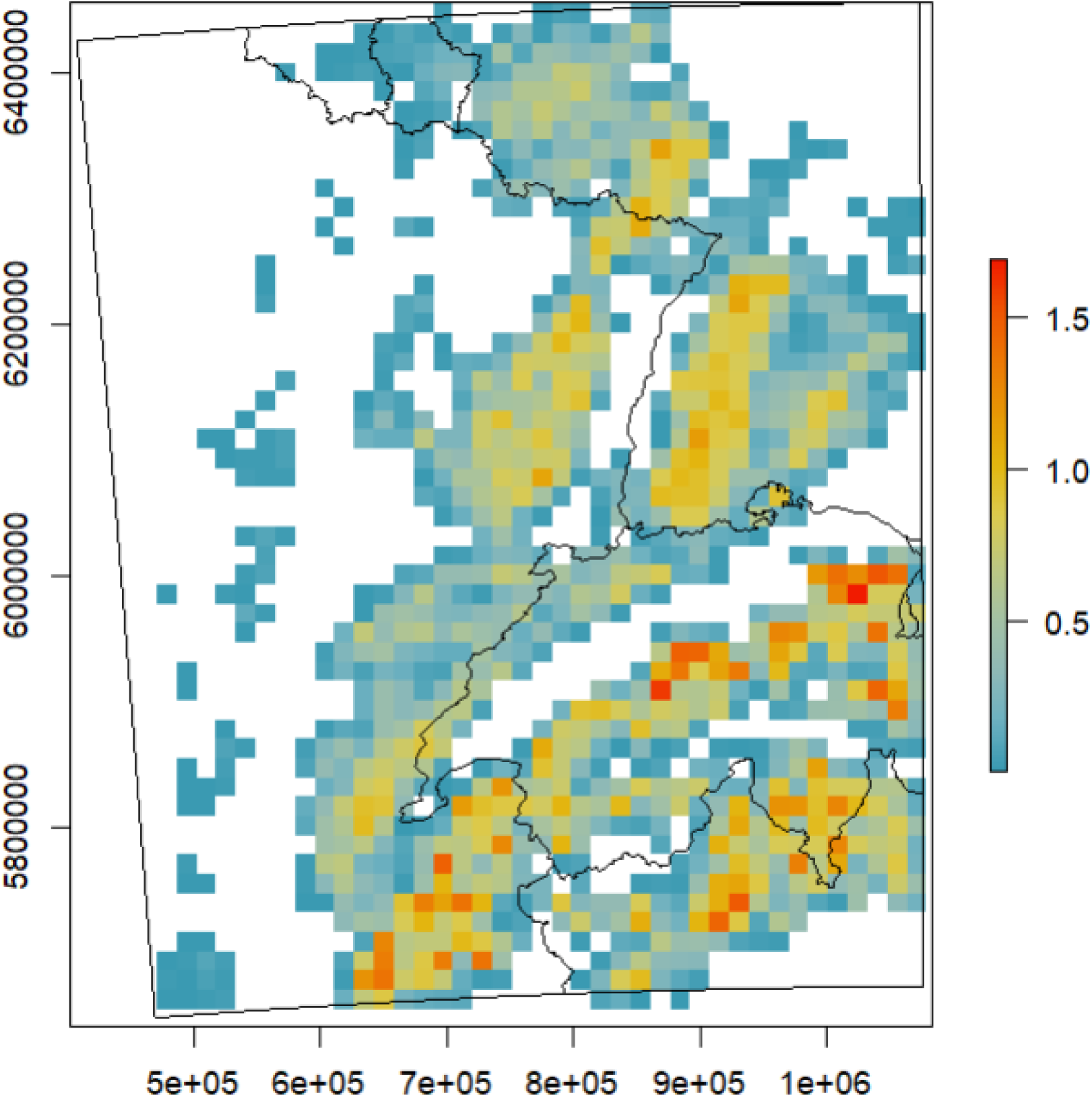
Adult resident lynx density per 100 km^2^ in the last year of simulation. Lynx density (color scale blue-yellow-red) are mean values over the 100 simulation replicates. White cells had a density equals to 0. Axis (x and y) are in meters from the WGS 84 / Pseudo-Mercator coordinate system.

## Discussion

### The well-established Alpine and Jura populations

The model predicted a steady growth for the Alpine and Jura populations, quickly reaching a more stable phase, indicating that they may soon be at saturation, especially for the Jura population. However, saturation could be higher than what we found in our simulations, depending on the strength of density-dependence processes (Zimmermann, Breitenmoser-Würsten, & Breitenmoser, 2007). Although our model assumed that the number of individuals an area can support is mainly defined by female territory size, it does not include a relationship between lynx behavior and density. Yet, studies found that lynx density also influences home range sizes (Pesenti & Zimmermann, 2013), and differently for females and males (Aronsson et al., 2016). For instance, in the North-Western Swiss Alps, lynx territories were much larger in the 80s when lynx density was lower compared to the 90s (Breitenmoser-Würsten et al., 2001). Simulated lynx density in 50 years was a little smaller than those estimated in the field for both populations (Gimenez et al., 2019; Pesenti & Zimmermann, 2013). Our model did not seem to allow a significant increase in lynx density. The Alpine and Jura areas could therefore support more individuals if such density-dependent mechanisms would occur.

### The Jura population as the crossroad of Western Europe lynx movements

The Jura population was found to be the only one connected by lynx exchanges with all the other populations which is coherent with observed data, also in terms of the number of individuals moving per year (Drouet-Hoguet et al., 2021). Individual exchanges simulated were low, with only a couple of individuals moving from their native population to establish their territory in another one on average per year. Exchanges were predicted to be the highest for the individuals born in the Alps and moving to settle down in the Jura Mountains, with two to three individuals per year after 25 years of simulation. In Switzerland, only a few lynx movements between the Alpine and Jura populations have been observed despite a large camera-trapping effort over the years. But even though camera trapping can detect movements between populations, it cannot guarantee an exhaustive sampling nor ensure that individuals successfully established in their destination, which contrasts with the computed results. However, Alpine and Jura populations also differ genetically, suggesting very few exchanges between them (Breitenmoser-Würsten & Obexer-Ruff, 2003). Our model may slightly over-estimate these dispersal movements due to the absence of movement barriers in the “habitat layer”.

When inspecting model output maps (Figs. 5 and 6), individuals are contained in the Swiss Alps, restricted to an area of suitable habitat almost totally surrounded by less favorable habitats (Fig. A.2). However, individuals have recently started to settle permanently on the Swiss Plateau (between the Alps and the Jura Mountains) predicted with less favorable habitats. Some have even reproduced successfully (F. Zimmerman, pers. comm.), indicating potential connections between the two populations in the future. On the other hand, our model suggests connectivity between the Jura and Alpine populations on the French side with continuity of a few forested corridors until the Chartreuse Mountains in the Alps (Zimmermann & Breitenmoser, 2007). Many observations were made via camera traps of lynx moving between the southern part of the Jura population and the Chartreuse Mountains (Bailly, 2021). However, movement barriers (e.g., highways) may prevent connections with the rest of the French Alps and the Alpine populations in Switzerland. Illegal killings, notably in Switzerland in movement corridors as well as a slow demographic dynamic of the Swiss population may also have prevented a larger development of the lynx population in the French Alps (Arpin et al., 2024).

Lynx movements simulated from the Jura to establish within the Vosges-Palatinian population were possible but rare. This is coherent with field monitoring that highlighted the presence of one male in the south of the Vosges Mountains, coming from the Jura Mountains during the winter 2014-2015 (Chenesseau & Briaudet, 2016; Hurstel & Laurent, 2016). Connectivity between these two populations remains far from optimal because of multiple barriers (e.g., highways, railroads and rivers) and restricted habitat that impede lynx dispersal between the two mountain ranges and increase collision risk (Zimmermann & Breitenmoser, 2007; Morand, 2016).

Regarding the Black Forest (including the adjoint Swabian Alb area), six males are known to have immigrated from the Swiss side of the Jura population since 2013 (unpublished data, Forest Research Institute Baden-Wuerttemberg; Stiftung KORA, 2017; Drouet-Hoguet et al., 2021), as well as two males from the Northeastern Alps (Herdtfelder, Schraml, & Suchant, 2021). Simulated lynx movements towards the Black Forest seem therefore coherent with field data. After 25 years of simulation, it is the Vosges-Palatinian population which seemed the most connected to the Black Forest. This is concurrent with the growth of the Vosges-Palatinian population, which could act as a source population only after several years, as it started only with a few individuals. It lasted several years until the first lynx immigrated to the Black Forest from the other populations in our simulation compared to the others. This is due to male establishment which is driven by female presence in our model, and because the Black Forest population did not have any female at first, new males arriving could not establish and were doomed to die while searching for females. A more realistic rule for male establishments would be to let them search for females but still allow them to settle after a defined period, even without females. It happens quite regularly that males show territorial behavior even without females (M. Herdtfelder, pers. Comm.).

### The increasing Vosges-Palatinian and Black Forest populations

Projections for the Vosges-Palatinian and Black Forest populations showed positive growth over the next 50 years. However, there was a difference in their initial populations. The lynx reintroduction program in the Palatinate Forest conducted since 2016 constituted a new larger initial population for the Vosges-Palatinian one (Scheid, Germain, & Schwoerer, 2021; Schwoerer, 2021) compared to the few individuals, only males (Drouet-Hoguet et al., 2021), located in the Black Forest. Growth rates reached higher values for the Black Forest population, but growth rates were very heterogeneous along the simulated period and confidence intervals around the mean were very large. Demographic stochasticity greatly impacted the Black Forest population because of its small initial population. Precision, as well as robustness, to interpret the results for this population were indeed lower than for the Vosges-Palatinian population. In the field, one female was reintroduced in the Black Forest by the end of 2023 to support this only-male population, but it died in Summer 2024 (www.baden-wuerttemberg.de), highlighting the high stochasticity of the recolonization of this habitat patch.

Exchanges of individuals between the Vosges-Palatinian population and the Black Forest one increased only at the end of the simulation, probably when both populations had more individuals. The two also seemed connected with the Jura population, highlighting the Upper Rhine meta-population structure. However, functional connectivity within the meta-population may not be at its best and may be altered by anthropogenic barriers. For example, in the Vosges-Palatinian population, at the *Col de Saverne,* this forest bottleneck is fragmented by both a highway and high-speed railway (Klar, Hermann, & Kramer-Schadt, 2006; Morand, 2016; Scheid, Germain, & Schwoerer, 2021). However, movements are possible as lynx from the Palatinate Forest are known to have crossed this pass, some even going back and forth (Idelberger et al., 2021; Scheid, Germain, & Schwoerer, 2021). Moreover, no female lynx has been observed until now dispersing from the Jura Mountains to the Black Forest crossing the Rhine valley that separates the Black Forest population from the Jura one, probably due to their conservative dispersal behavior compared to males (Port et al., 2021). Monitoring data from 2021 indicate that one female from the north-eastern Swiss population crossed this barrier for the first time (M. Herdtfelder, pers. comm.).

### Model limitations

Although predicted growth rates did not seem very sensitive to model calibration and model predictions were well validated (Appendices C and D), several aspects of our IBM could still be improved.

The “habitat layer” is defined in categories with strict “barriers” that the simulated lynx cannot cross, whereas the species may be tolerant of human activities (Basille et al., 2009; Bouyer et al., 2015). Lynx may live near urban areas, cross small lakes (F. Zimmermann, pers. comm.), and sometimes large rivers. In that context, the “habitat layer” could be improved by being defined as a continuous variable representing a degree of selection or permeability of the landscape to avoid prohibiting the movement through certain landscape elements. The “habitat layer” could also be improved by accounting for roads and their associated structures as movement barriers (e.g., highways and other types of roads difficult to cross, over-pass facilitating the movement) instead of only mortality sources (Klar, Hermann, & Kramer-Schadt, 2006; Marchand et al., 2017).

In our model, lynx movement is not impeded by roads. Including permeability of these linear barriers to the lynx movements may help to refine the behavior rules (Marchand et al., 2017). We could also redefine the movement behavior rules to be sex or age-dependent. Indeed, females seem more conservative (i.e., disperse close to their natal range) compared to males and some males can disperse over long distances (Port et al., 2021). Moreover, lynx movement and therefore the impact of habitats and collision risk were modeled only for dispersing individuals. Movements inside territories were not simulated for resident lynx and we assumed an even use of the space. Model predictions could be different if resident individuals would move across their territory in an uneven way, increasing or decreasing the averaged collision risk of their home range. However, mortality risk for residents, thanks to their knowledge about their territory where and how to cross linear infrastructures, would probably be lower than for dispersing individuals.

Moreover, our model predictions were made in a static environment, as the landscape did not change over the simulated period. Modifying the landscape (e.g., agricultural intensification, forest cover increase) would give different lynx population predictions. However a landscape model would be necessary to predict a realistic future landscape, or different scenarios could be defined (Bauduin et al., 2018). We used a medium-term simulation period of 50 years in which a static landscape was a reasonable assumption.

Finally, we did not include any genetic aspect in the demographic rules of our SE-IBM (Mueller et al., 2022). A collective expertise recently done in France on lynx viability showed that inbreeding depression could greatly impact population size and extinction risk (Arpin et al., 2024). We restricted our predictions over a 50-year period to limit the (non-included) effects of inbreeding depression on population dynamics. However, even within this time frame, population trends could shift from increasing when inbreeding depression is excluded, to potentially decreasing when included. Therefore, our results need to be interpreted carefully and more in lights of population relative differences than raw values for each population *per se*.

### Perspectives for assessing lynx conservation strategies

Our model used to evaluate lynx population persistence could be applied to better understand aspects of lynx conservation that we did not include yet, such as illegal killings (Heurich et al., 2018). Moreover, thanks to its individual-based structure, a genetic component could easily be included to track relatedness between individuals and allow studying inbreeding risk and Allee effects (Arpin et al., 2024; Premier et al., 2020). Our model could also be used to test the effect of different scenarios, either by modifying the lynx population sizes (e.g., due to illegal killings or reintroduction programs) or the landscape at different scales (e.g., green bridges, roads construction, habitat destruction or restoration) and assess the potential benefits or negative effects on lynx population persistence. Modifications of road networks to improve connectivity, such as the removal of road segments or the addition of overpasses or underpasses could be tested. Then, their effect could be included through an updated layer of collision probabilities and the population persistence calculated accordingly. Similarly, other modifications of land cover (e.g., restoring forest areas) could also be tested. This model could be very useful for stakeholders working on corridors and the reduction of lynx-vehicle collisions as well as reintroduction programs and species acceptance. Helping reducing road mortality using model simulations is one of the objectives of the French national action plan (Gatti, 2022).

Using IBMs, Herdtfelder (2012) showed that Black Forest population reinforcement with lynx females might be one solution considering that habitat is of good quality in this area. The collective expertise on lynx viability also used a model similar to the SE-IBM presented here to make recommendations on lynx translocations in the Vosges area to support the population viability (Arpin et al. 2024). IBMs are great tools to evaluate the impact of such management actions. However, because our model does not include human attitudes towards the species, we warn against its blind use to assess the effect of reinforcement on lynx long-term persistence without accounting for the human dimension component. Illegal killings, as they occurred after the 90’s reintroduction program in the southern part of the Vosges Mountains (Vandel et al., 2006), as well as more recent ones (Germain, 2020), may be an additional mortality that the model does not account for. The extent of acceptance towards species reinforcement from some local stakeholders is an essential element (Charbonnel & Germain, 2020). In this case, we recommend extending our model to include the dynamics of the whole socio-ecosystem (Behr, Ozgul, & Cozzi, 2017; Guerrero et al., 2018).

## Conclusion

In this paper, we built and analyzed a spatially-explicit individual-based model to forecast the fate of lynx populations over the next 50 years. Our results suggest that exchanges of individuals between Vosges-Palatinian, Black Forest, Jura and Alpine populations to establish new territories were limited, and emphasize that the Jura population plays the role of a Western Europe crossroad. Overall, lynx persistence in the Upper Rhine meta-population and the Alpine population over the next 50 years seems likely on a large scale by considering road mortality and habitat selection.

## Data sharing

The lynx SE-IBM and all the necessary layers to run it (Appendix A), as well as the simulation outputs and the code to analyze those are available on https://github.com/SarahBauduin/appendix_lynxIBM.

## Acknowledgements

We thank all the volunteers from the “Réseau Loup-Lynx” who collected data on the field. SB and OG were funded by French National Research Agency (ANR-16-CE02-0007). SB was funded as well by OFB and OG was funded by CNRS and the “Mission pour l’Interdisciplinarité” through the “Osez l’Interdisciplinarité” initiative. CEFE, Cerema and CROC were funded by CILB, MTES (ITTECOP) and FRB through the research program ERC-Lynx. Cerema received support from the “ Direction générale des infrastructures de transports et de la mer (Ministère de la transition écologique)”. CROC was funded in 2019/2020 by the European Union within the framework of the Operational Program FEDER-FSE “Lorraine et Massif des Vosges 2014– 2020”, the “Commissariat à l’Aménagement du Massif des Vosges” for the FNADT (“Fonds National d’Aménagement et de Développement du Territoire”), the DREAL Grand Est (“Direction Régionale pour l’Environnement, l’Aménagement et le Logement”), the “Région Grand Est” and the “Fondation d’entreprise UEM”. This research is partly a product of the DISCAR group funded by the synthesis centre CESAB of the French Foundation for Research on Biodiversity. We thank OFB and Réseau Loup-Lynx for the lynx presence and lynx-vehicle collision data to build the habitat and collision models; OFB, Réseau Loup-Lynx, KORA and Bundesamt für Naturschutz for population data to create the initial populations; OFB, Réseau Loup-Lynx and KORA for the collision data for model validation; and KORA and Stiftung Natur und Umwelt Rheinland-Pfalz for the female telemetry data for model validation. We thank Elodie Vercken, Hector Ruiz and Henrik Andren for their very constructive reviews. A preprint version of this article has been peer-reviewed and recommended by PCIEcology (https://doi.org/10.24072/pci.ecology.100408).

### Appendix A Detail of the lynx spatially-explicit individual-based model

A complete description of the model following the Overview, Design concepts, and Details (ODD) protocol (Grimm et al., 2006, 2010) is provided in Appendix B.

Our lynx spatially-explicit individual-based model is made of four components (main text, Fig. 2). The first component represents the impact of road network on lynx survival via predicted collision probabilities. The second component represents the impact of land cover on lynx space use with the definition of different lynx habitat types. These first two components are spatial layers influencing the behavioral rules followed by simulated lynx individuals. The third component represents the initial lynx populations made of lynx individual’s locations and characteristics used to launch the SE-IBM. The fourth component details all SE-IBM rules including lynx demography, dispersal movement and establishment.

#### Impact of road network (SE-IBM 1^st^ component)

We first built a risk model to predict collision probabilities between lynx and vehicles within the 1 km² resolution gridded study area. We used lynx mortality events recorded by the wolf-lynx monitoring framework implemented in France (Duchamp et al., 2012, Réseau Loup-lynx https://www.loupfrance.fr/suivi-du-loup/reseau-loup-lynx/), which represents 84 collision events recorded between 1982 and 2016, to train a logistic regression explaining lynx collisions using lynx presence and both road and environmental characteristics (Visintin, van der Ree, & Mccarthy, 2017; Visintin et al., 2018) (Table A.1). We used the IGN route500 ^©^ for France (IGN ROUTE 500, 2018) and OpenStreetMap^©^ for the other countries (Geofabrik OpenStreetMap, 2014) as data sources to extract total road length per cell and the type of roads of the longest road segment in each cell. We classified the road segments as “highways” (i.e., “Type autoroutier” in route500 ^©^ data, “motorway”, “motorway_link”, “trunk” and “trunk_link” in OpenStreeMap data), “main road” (i.e., “Liaison principale” in route500 ^©^ data, “primary” and “primary_link” in OpenStreeMap data), “secondary road” (i.e., “Liaison régionale” in route500 ^©^ data, and “secondary” and “secondary_link” in OpenStreeMap data) and “local road” (i.e., “Liaison locale” in route500 ^©^ data, “tertiary”, “tertiary_link” and “unclassified” in OpenStreeMap data). For environmental characteristics, we used Corine Land Cover^©^ at 100 m of resolution for all Europe (Copernicus, 2012) to calculate the proportion of urban area in each cell (i.e., human presence).

The best model identified included the total road length, the type of road of the longest road segment and the proportion of urban area (Table A.2). Distance to highways and human density were also tested to explain lynx-vehicles collisions but were not significant. We used this model to predict a collision probability, ranging between zero and one in each cell of our study area, to create the “collision layer” (Fig. A.1). Lynx presence was also included as explanatory variable in the model. To predict collision probabilities, we defined the lynx as present everywhere on the gridded study area because simulated individuals in the SE-IBM suffer from collision probability on the cells they are located so the mortality value should already account for the lynx presence. Study area grid cells without road intersecting them have a zero-collision probability.

**Table A.1:** Statistical details for the non-categorical variables included in the best model identified to explain lynx-vehicle collisions.

| Variable | Unit | Mean | Standard deviation | Minimum value | Maximum value |
| --- | --- | --- | --- | --- | --- |
| Lynx presence | Years of presence | 14.5 | 12.0 | 0 | 35 |
| Total road length | Meters | 1212 | 728 | 0 | 8084 |
| Urban area | Percentage | 7.9 | 16.9 | 0 | 100 |

**Table A.2:** Parameter estimates for the best model identified to explain lynx-vehicle collisions (logistic regression parameters).

| Parameter | Estimate | Standard error | Z value | P-value |
| --- | --- | --- | --- | --- |
| Intercept | -6.29204 | 0.24903 | -25.267 | < 2e-16 |
| Lynx presence | 1.11454 | 0.15664 | 7.115 | 1.12e-12 |
| Total road length | 0.34289 | 0.10107 | 3.392 | 0.000693 |
| Longest segment<br>== highway | 1.69306 | 0.53300 | 3.176 | 0.001491 |
| Longest segment<br>== main road | 1.96672 | 0.27082 | 7.262 | 3.81e-13 |
| Longest segment<br>== secondary road | 0.53213 | 0.31437 | 1.693 | 0.090517 |
| Urban area | -0.41059 | 0.16709 | -2.457 | 0.013996 |

**Figure A.1:**
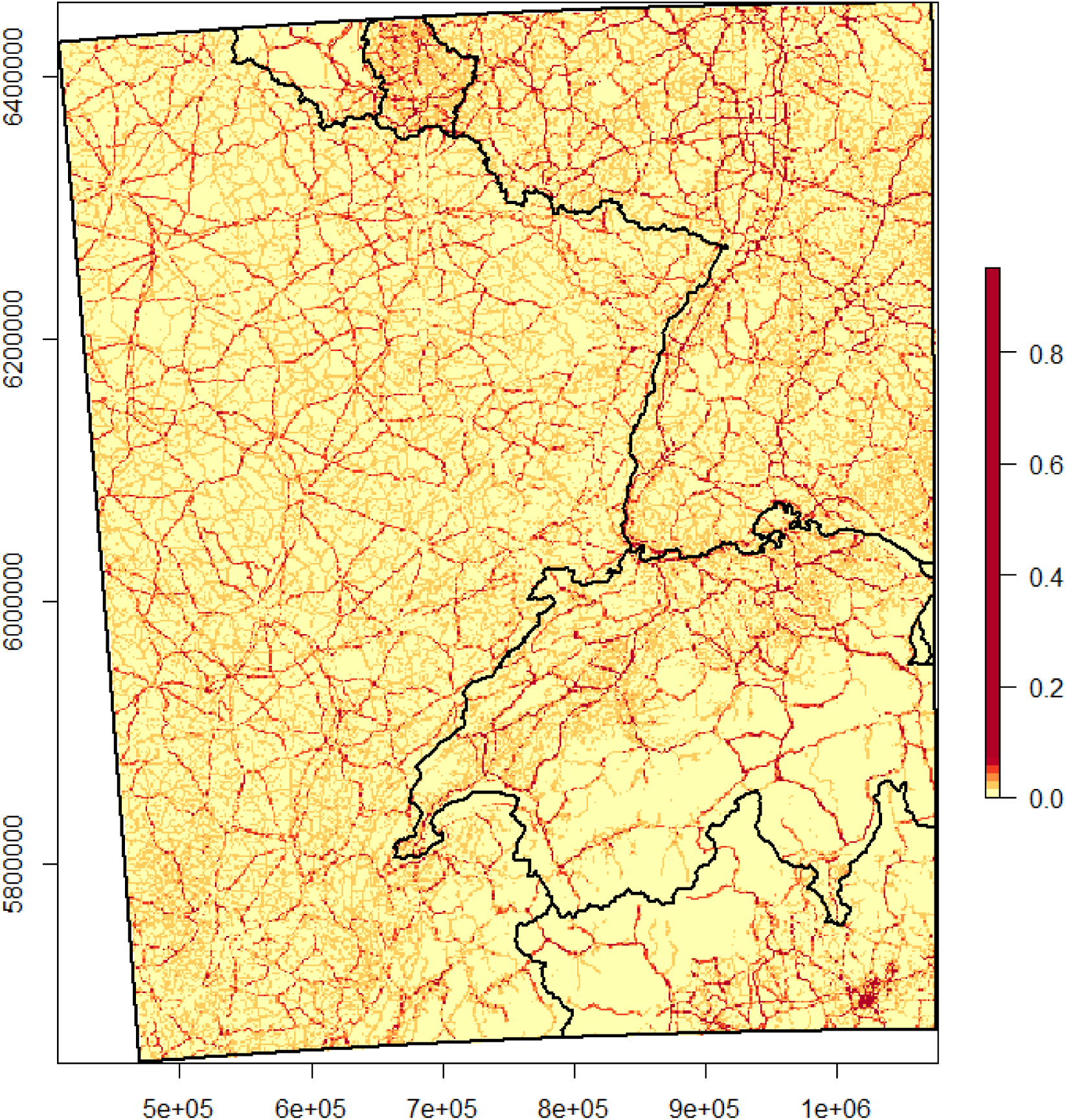
“Collision layer” with collision probabilities between lynx and vehicles estimated between zero and one (yellow to red color scale). Collision probabilities are assigned to each cell of the gridded landscape over the whole study area (black rectangle) used in the lynx SE-IBM using the collision model. Limits of the countries (main text, Fig. 1) are overlaid in black over the map.

#### Impact of land cover (SE-IBM 2^nd^ component)

We built a habitat model to define a habitat type for each cell within the 1 km² resolution gridded study area. We used a multi-year occupancy model (Isaac et al., 2014; Outhwaite et al., 2018) in a Bayesian framework to explain regular lynx presence using land cover types important to lynx (agricultural fields, forest and open land), the distance to highways (Basille et al., 2013), and human density (Table A.3). The model was calibrated using French data for the lynx presence from 1994 to 2017 (data: Réseau Loup-Lynx https://www.loupfrance.fr/suivi-du-loup/reseau-loup-lynx/). The model used both lynx detection and non-detection data over the surveyed area each year (https://ecologicalstatistics.shinyapps.io/flexdashboard_effort/). We used Corine Land Cover^©^ at 100 m of resolution for all Europe (Copernicus, 2012) to calculate the proportion of agricultural and cultivated fields, forest, pasture and open land in each cell. We used the IGN route500 ^©^ for France (IGN ROUTE 500, 2018) and OpenStreetMap^©^ for the other countries (Geofabrik OpenStreetMap, 2014) as data sources to calculate the distance from each cell center to the nearest highway. We also used IGN^©^ data for France (IGN ADMIN-EXPRESS-COG, 2018) and NASA^©^ data for the other countries (CIESIN GPWv4, 2015) to calculate the mean human density per cell on a log scale. Finally, we also added a yearly fixed effect and a random site effect to the model. The best model identified to explain lynx presence included presence of agricultural fields, forests, and open lands, distance to highways and human density. Parameters values (Table A.4) were estimated for the occupancy on the logit scale. Shrub cover and road length were also tested to explain lynx presence but were not significant. We used the inverse logit function to predict lynx occupancy probability in our entire gridded study area for the last year (i.e., 2017). We then categorized the cells to obtain a “habitat layer” with the four habitat types defined by the habitat categorization of Kramer-Schadt et al. (2004): “breeding”, “dispersal”, “matrix” and “barrier” (Fig. A.2). All cells where lynx occupancy was predicted as non-null (above or equal a threshold of 0.01) were considered “breeding” habitat. All forested areas not already “breeding” habitat were considered as “dispersal” habitat. Cells with more than half of their surface covered by water or urban area defined by Corine Land Cover^©^ (Copernicus, 2012) were considered “barrier” for lynx movements. The rest was considered “matrix” area (i.e., habitat not favorable for lynx but that can be traversed by dispersers). The map of categorized cells for our study area defined the “habitat layer” used in the SE-IBM (Fig. A.2).

**Table A.3:**
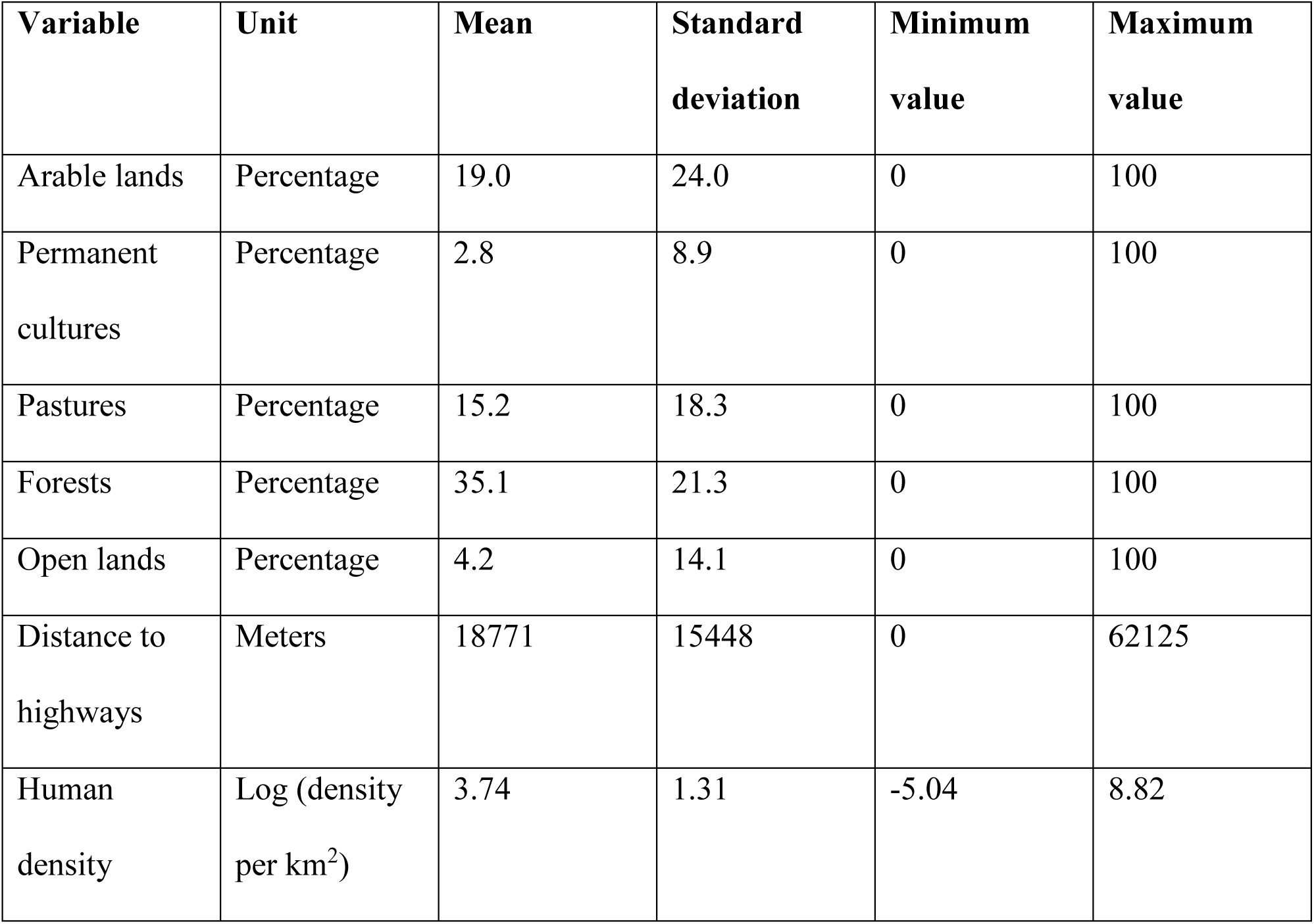
Statistical details for the variables included in the best model identified to explain lynx presence.

| **Variable** | **Unit** | **Mean** | **Standard deviation** | **Minimum value** | **Maximum value** |
| --- | --- | --- | --- | --- | --- |
| Arable lands | Percentage | 19.0 | 24.0 | 0 | 100 |
| Permanent cultures | Percentage | 2.8 | 8.9 | 0 | 100 |
| Pastures | Percentage | 15.2 | 18.3 | 0 | 100 |
| Forests | Percentage | 35.1 | 21.3 | 0 | 100 |
| Open lands | Percentage | 4.2 | 14.1 | 0 | 100 |
| Distance to highways | Meters | 18771 | 15448 | 0 | 62125 |
| Human density | Log (density per km<sup>2</sup>) | 3.74 | 1.31 | -5.04 | 8.82 |

**Table A.4:** Parameter estimates for the best model identified to explain lynx presence (Bayesian occupancy model parameters).

| **Parameter** | **Mean** | **Standard deviation** |
| --- | --- | --- |
| Arable lands | -1.06 | 0.23 |
| Permanent cultures | -0.85 | 0.24 |
| Pastures | 0.68 | 0.15 |
| Forests | 1.67 | 0.16 |
| Open lands | 0.49 | 0.17 |
| Distance to highways | -0.70 | 0.14 |
| Human density | 0.77 | 0.15 |
| Yearly fixed effect (estimate for the last year) | -6.57 | 0.27 |

**Figure A.2:**
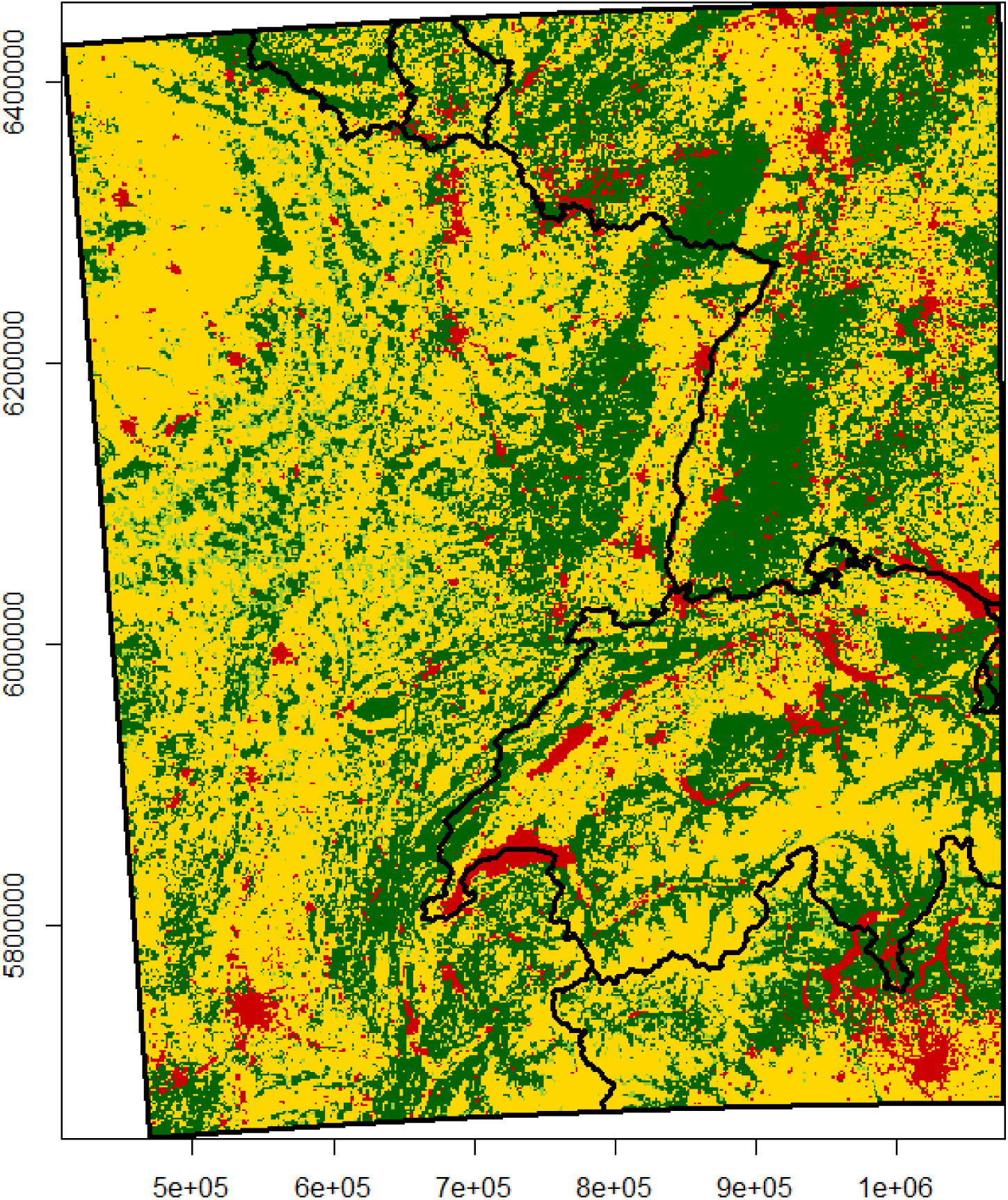
“Habitat layer” with lynx habitat types as breeding habitat (dark green), dispersal habitat (light green), matrix (yellow) and barrier (red). Habitat types are assigned to each cell of the gridded landscape over the whole study area (black rectangle) used in the lynx SE-IBM using the habitat model. Limits of the countries (main text, Fig. 1) are overlaid in black over the map.

#### Lynx initial populations (SE-IBM 3^rd^ component)

The initial populations to launch the SE-IBM were built with generated locations and characteristics for the lynx populations using the best data available at the time of the study (Fig. A.3). We used cells of regular lynx presence for France (for the time period of 01/04/2013 to 31/03/2017 from the Réseau Loup-Lynx, main text, Fig. 1), Switzerland (for the time period of 01/01/2015 to 31/12/2017 from the KORA, main text, Fig. 1) and Germany (for the time period of 01/05/2017 to 30/04/2018 from the Bundesamt für Naturschutz, main text, Fig. 1). In these cells, we extracted “breeding” and “dispersal” areas and sampled random lynx locations in these areas to locate individuals within the different populations. In France, we used the most reliable lynx density estimate (1.14 lynx per 100 km^2^; Gatti et al., 2014), regardless of differences in local densities (Gimenez et al., 2019), to convert the area of regular presence into a number of individuals to create. Ninety-two individuals were generated and dispatched over “breeding” and “dispersing” areas in France: in the Vosges Mountains (500 km^2^ of presence), in the French Jura (7,700 km^2^ of presence) and in the French Alps (500 km^2^ of presence). A density of 1.14 lynx per 100 km^2^ may be an over- or an under-estimate of the lynx density in certain areas but in the absence of local density for each French population, we used this mean value. On average, this method led to 4.5 individuals in the French Alps (sd = 2.1), 77.0 individuals in the French Jura (sd = 3.4), and 8.4 individuals in the Vosges Mountains (sd = 2.7). In Switzerland, we generated 230 individuals distributed in the different areas of presence according to estimated local population sizes in 2017 (data: F. Zimmermann, pers. comm.). In Germany, 11 lynx (i.e., reintroduced lynx still alive and their offsprings) were identified in the Palatinate at the end of April 2018 (Scheid, Germain, & Schwoerer, 2021) and four male lynx were identified in the Black Forest (Wölfl et al., 2021). Therefore, we generated 11 lynx in Palatinate Forest and four lynx in the Black Forest area.

Except for reintroduced individuals for which we knew their characteristics, we randomly assigned individuals’ sex (male or female) according to the ratio 1:1 usually observed at birth (Breitenmoser et al., 1993; Jedrzejewski et al., 1996). We also randomly assigned an age between 2 and 15 as lynx usually live until 15-17 years old (e.g., Breitenmoser-Würsten, Vandel, et al., 2007). Age is defined as zero for the first year of life, one during the second year of life, etc. For the Palatinate part of the Vosges-Palatinian population, we assigned known age and sex (five females and six males). In the Vosges Mountains, only males were detected in 2017 thanks to the camera trap survey design implemented (Charbonnel & Germain, 2020) and no case of reproduction was reported during the years preceding our analyzes. We then only defined males for the individuals located in the Vosges part of the Vosges-Palatinian population, with ages randomly generated. We also only defined males for the Black Forest population. All generated individuals were defined as “disperser” to avoid the bias of defining territories by ourselves. They find their territories on their own as defined by the SE-IBM rules.

We also used lynx regular presence data to define the “population layer” (Fig. A.3). This layer took the gridded study area where each cell was assigned one of the four lynx populations (main text, Fig. 1), taking the population from the closest lynx regular presence cell. This “population layer” delineates population areas. It is used to define in which population kitten were born and identified individual exchanges between populations.

**Figure A.3:**
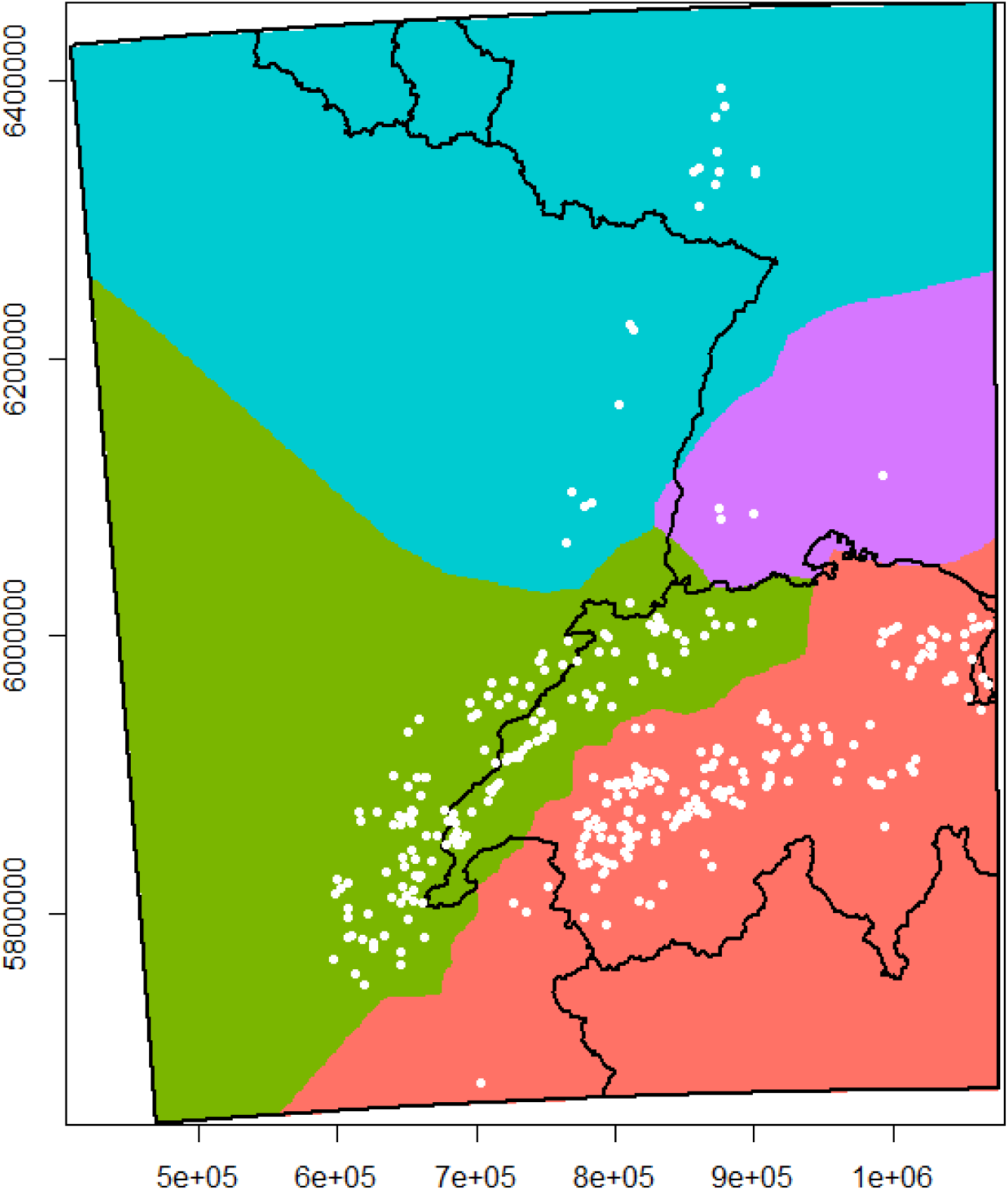
“Population layer” with the example of an initial population over the whole study area (black rectangle) used in the lynx SE-IBM using the habitat model. Limits of the countries (main text, Fig. 1) are overlaid in black over the map. Areas for the Vosges-Palatinian (blue), Black Forest (purple), Jura (green) and Alpine (red) populations were defined using the cells of lynx regular presence as in 2017-2018. Each cell of the gridded study area was assigned the population from its closest cell of lynx regular presence. White dots represent one initial population (i.e., simulated lynx released at the beginning of one simulation replicate) generated using the cells of lynx regular presence, the density or number of lynx in each population and the “habitat layer” to place these theoretical individuals in breeding or dispersal habitats. A different initial population was used at each simulation replicate; we used the same cells of lynx regular presence, the same density or number of lynx in each population and the same “habitat layer”, only the generated locations, and chosen sex and age for unknown individuals, were different.

#### SE-IBM rules according to lynx ecology (SE-IBM 4^th^ component)

Lynx individuals are simulated over a landscape represented as a grid of 1 km² resolution, covering the whole study area and encompassing the four lynx populations (main text, Fig. 1). The gridded study area resolution corresponds to lynx’s perceptual range (Haller & Breitenmoser, 1986) as well as to the resolution of previous lynx IBMs (Kramer-Schadt et al., 2004, 2011; Kramer-Schadt, Revilla, & Wiegand, 2005). Two variables characterize the gridded study area: a probability of lynx-vehicle collision between zero and one (“Collision layer”, Fig. A.1) and a habitat type among “breeding”, “dispersal”, “matrix”, and “barrier” (“Habitat layer”, Fig. A.2). Simulated individuals are characterized by their “disperser” (i.e., not established on a territory and in search of one) or “resident” status (i.e., established on a defined territory), their age and their sex.

Simulated resident individuals follow rules on a yearly time step (Fig. A.4). They do not move (i.e., their movements inside their territory are not simulated), they hold a territory and they may reproduce once a year. They suffer two types of annual mortality: a fixed baseline mortality and a spatial one derived from the “collision layer”. Simulated dispersing lynx follow rules on a daily time step (Fig. A.4). They do not have a territory yet and move every day along the gridded study area, searching for a place to establish themselves. Their dispersal movements (Fig. A.5) and search for a territory (Fig. A.6) are driven by the “habitat layer”. At each step, individuals can die from the spatial mortality derived from the “collision layer”, and daily from a fixed baseline mortality.

**Figure A.4:**
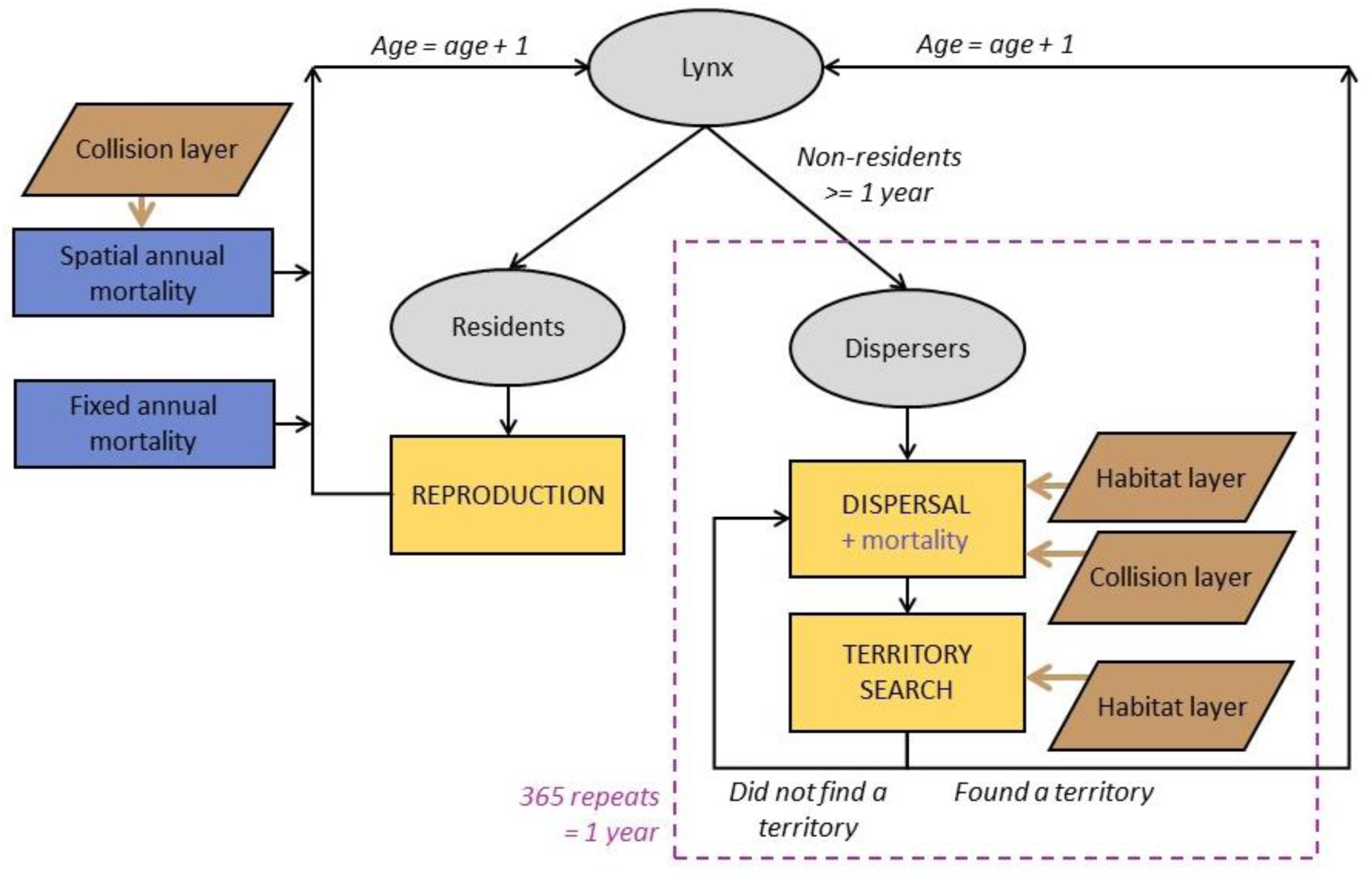
Diagram of the main structure of the SE-IBM with the main events affecting both resident and dispersing simulated lynx. Grey circles represent individuals. Yellow boxes represent SE-IBM main events, detailed in model description. Blue boxes or blue writing represent different mortality causes individuals suffer. Brown boxes represent environmental layers used in the SE-IBM and brown arrows point where behavioral rules are constrained by these layers. Black arrows indicate the flow of the model. Dispersers can go through dispersal and territory search every day during a year, as long as they do not find a territory (Figs. A.5 and A.6).

The modeling cycle starts with the dispersing individuals. Dispersers are individuals of one year old and older that do not hold a territory yet. Both dispersing males and dispersing females move on the gridded study area, one cell at a time, from one to 45 times per day (Fig. A.5), following the same rules. The number of steps individuals move per day is sampled, each day for each disperser, from a non-linear distribution (Kramer-Schadt et al., 2004). Dispersers follow a correlated habitat-dependent walk in a two-step process: first, they favor their habitat preferences and then, maintain their previous direction with a certain probability (i.e., correlation factor; Kramer-Schadt et al., 2004). This movement process has been rigorously tested with inverse fitting, and “pattern-oriented modelling” (Kramer-Schadt et al., 2007) using telemetry data of six dispersing lynx (five females and one male) followed in the Swiss Jura Mountains between 1988 and 1991. First, dispersers choose in which habitat type they will move next. Dispersers favor “breeding” and “dispersal” habitats without distinction between the two when moving and tend to avoid “matrix” habitats. By contrast, they never use “barrier” habitats. The choice to move into the habitat “breeding/dispersal” or “matrix” depends on the types of their nine available cells for their next step (i.e., their eight surrounding cells plus the one they are currently on as they also can choose not to move). If the nine cells are all of one type, excluding the “barrier”, so they are either only “breeding/dispersal” (with or without “barrier”) or only “matrix” (with or without “barrier”), the only available habitat type is selected. If the available cells for an individual are a mix between “breeding/dispersal” and “matrix” habitats (with or without “barrier”), there is a probability of 0.03 times the number of “matrix” cells among the nine ones, to choose the “matrix” habitat for the next step (Kramer-Schadt et al., 2004). For example, if an individual has three “matrix” cells available, there is a probability of 0.09 that it will choose a “matrix” cell for its following location. Second, once the habitat type is selected, the choice of the particular cell to move on, among the ones of the selected habitat type, is given by the correlation part of the movement. Individuals follow a correlated movement with a probability equal to 0.5, except for the first step of the day where there is no correlated movement (Kramer-Schadt et al., 2004). If the movement is not correlated, the choice of the next cell among the ones of the selected habitat type is random. If the movement is correlated, the chosen cell is the one maintaining the most of the individual’s current direction among the ones of the selected habitat type. The chosen cell is then where the simulated lynx is moving to. Dispersers try to minimize their time spent in “matrix” habitat and they can use their memory to return to a previously visited “breeding/dispersal” habitat when needed. Lynx do not move more than nine cells inside “matrix” habitats, so if an individual already stepped nine consecutive times in “matrix” cells and the chosen cell for its next step is again of “matrix” type, it will use its memory and return to the last “breeding/dispersal” cell visited (Kramer-Schadt et al., 2004). Finally, dispersers rotate towards their chosen cell and move on their center. Once dispersers move to their next cell, they may die from the spatial mortality due to vehicle collision. This spatial mortality is the collision probability from the “collision layer” for the cell of their new location with a correction factor to transform the value into a per step collision mortality. If dispersers survive, they search for a territory. If found, they stop moving, establish a territory, and they will become resident the following year. Dispersers that do not find a territory to establish on their new location keep moving, as many steps during the day as simulated for them at the beginning of the day. At the end of the day, all individuals that dispersed during the current year may die from a fixed daily mortality probability (i.e., baseline mortality, Table B.1). Dispersers that have not established during the day can move and search for a territory every day during the year. The daily spatial mortality is also applied to individuals which dispersed during the year but already found a territory during previous days. Regarding spatial mortality, as movements inside territories are not simulated and they were not considered as resident at the beginning of the year, a per step spatial or annual mortality cannot be applied. A daily spatial mortality is then calculated as their future spatial mortality with their resident status divided by 365.

**Figure A.5:**
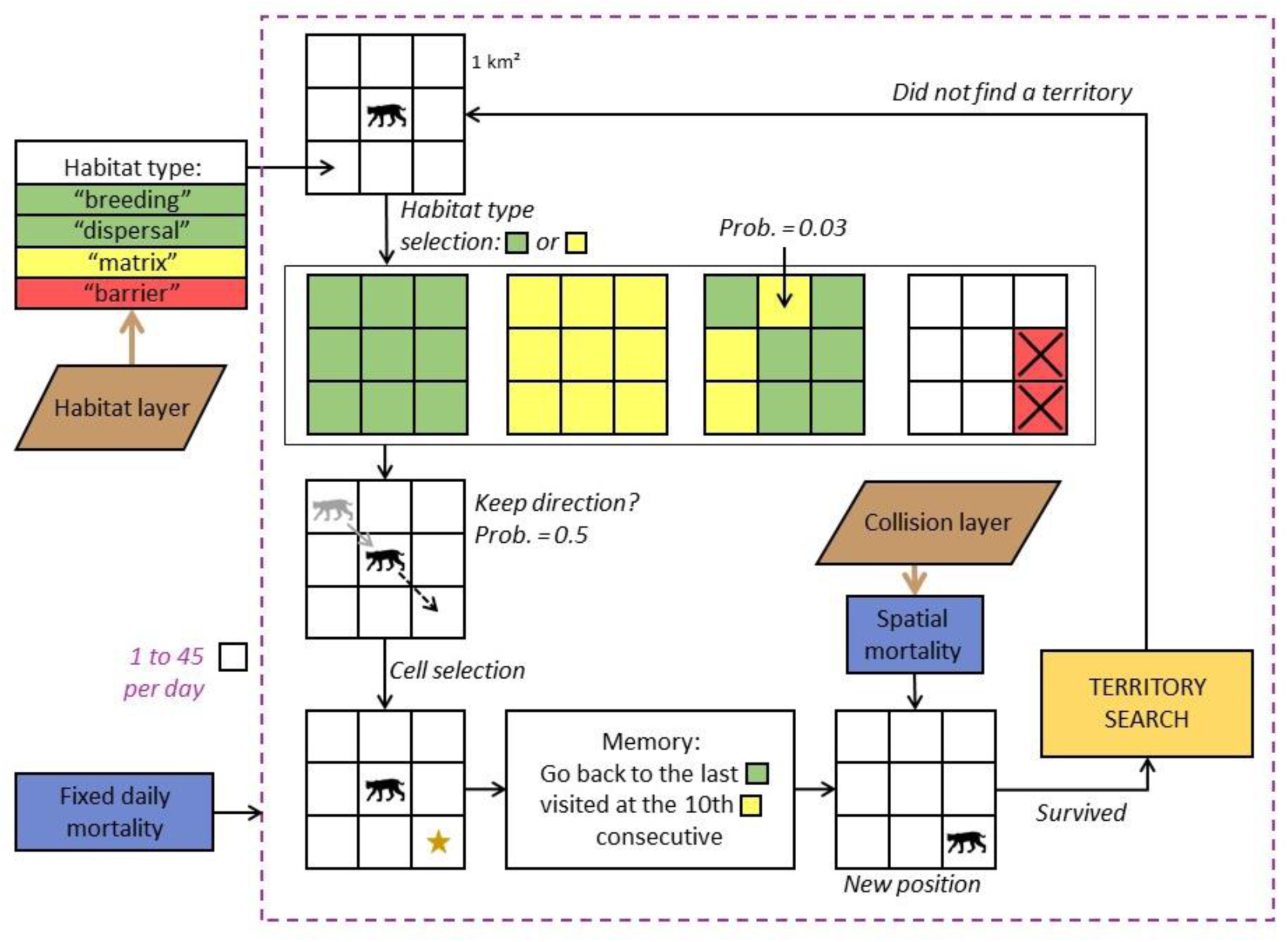
Diagram of simulated lynx dispersal movements in the lynx SE-IBM. The loop represents one step. Individuals move one cell at the time per step, up to 45 a day or until they find a territory to establish (yellow box). Individual movement is constrained by habitat types in their surroundings defined by “habitat layer” (brown box), probabilities to step into different habitat types and to keep a direction (i.e., correlated movement), and their memory. Blue boxes represent mortalities an individual suffers, a spatial one each time reaching a new location, derived from the “collision layer” (brown box) and a daily one.

Dispersing individuals arriving at a new location have a different strategy to search for a new territory to establish regarding their sex (Fig. A.6). Females mainly look for good habitat (which represent high prey availability) while males search for breeding opportunities and seek female presence (Breitenmoser-Würsten, Zimmermann, et al., 2007b). Female dispersers need to be on a cell of habitat type “breeding”, defined by the “habitat layer”, and with enough unoccupied “breeding” habitats around their position to establish their territory. Territories can have any shapes but they must only be composed of connected cells of “breeding” habitat (search in 8-cell directions). The size of the territory females needs to define is the number of connected “breeding” cells. It represents the female territory core area (Kramer-Schadt, Revilla, & Wiegand, 2005) and depends on the population area where they are located in. We defined the size of these territories to establish as the size of the 95% kernel ranges measured in the field as kernel are computed based on the most used habitats (i.e., “breeding habitat”). When a female builds a territory in the model, the cells used are no longer available for other females. Territory size for females in the Alpine population is equal to 76 km^2^ (Breitenmoser-Würsten et al., 2001) and to 119 km^2^ in the Jura population (Breitenmoser-Würsten, Zimmermann, et al., 2007b). As we did not have a reliable estimate for the Vosges-Palatinian and the Black Forest populations, we assigned them the value of the Jura population due to the similarity of the habitat structure (Fig. A.2). Once a female has established a territory, she becomes resident. If a resident male already established his territory nearby, the male includes the new resident female’s territory into his own, if he has less than three females already associated. Then, the two individuals may potentially reproduce the following year. That means, the male’s dispersal strategy to establish is female-dependent. At their new location, dispersing males check if they arrived inside the territory of a resident female available (i.e., with no male associated). If a male finds a female, he pairs with her and his territory becomes the same as the one from the female je joined. He also looks at nearby territories for up to three other available females to pair with. Males can pair with females whose territories are within the size of their maximum territories. The maximum distance between the male location and a female territory (i.e., a cell of her territory) is the radius of the 95% kernel ranges estimated for males. It is equal to 6.6 km for males in the Alpine population (95% kernel range = 137 km^2^; Breitenmoser-Würsten et al., 2001) and to 8.5 km for the other populations (95% kernel range = 226 km^2^; Breitenmoser-Würsten et al., 2007b). If there are several females available for a particular male, the closest one(s) are chosen. The male can potentially reproduce the following year with every three females. The male territory is defined by the union of all the female territories he has paired with. If a disperser cannot find a territory to create (for female) or to join (for male) at their location, it continues dispersing. If all the females a male is paired with die, the male start to disperse again in search of new available female.

**Figure A.6:**
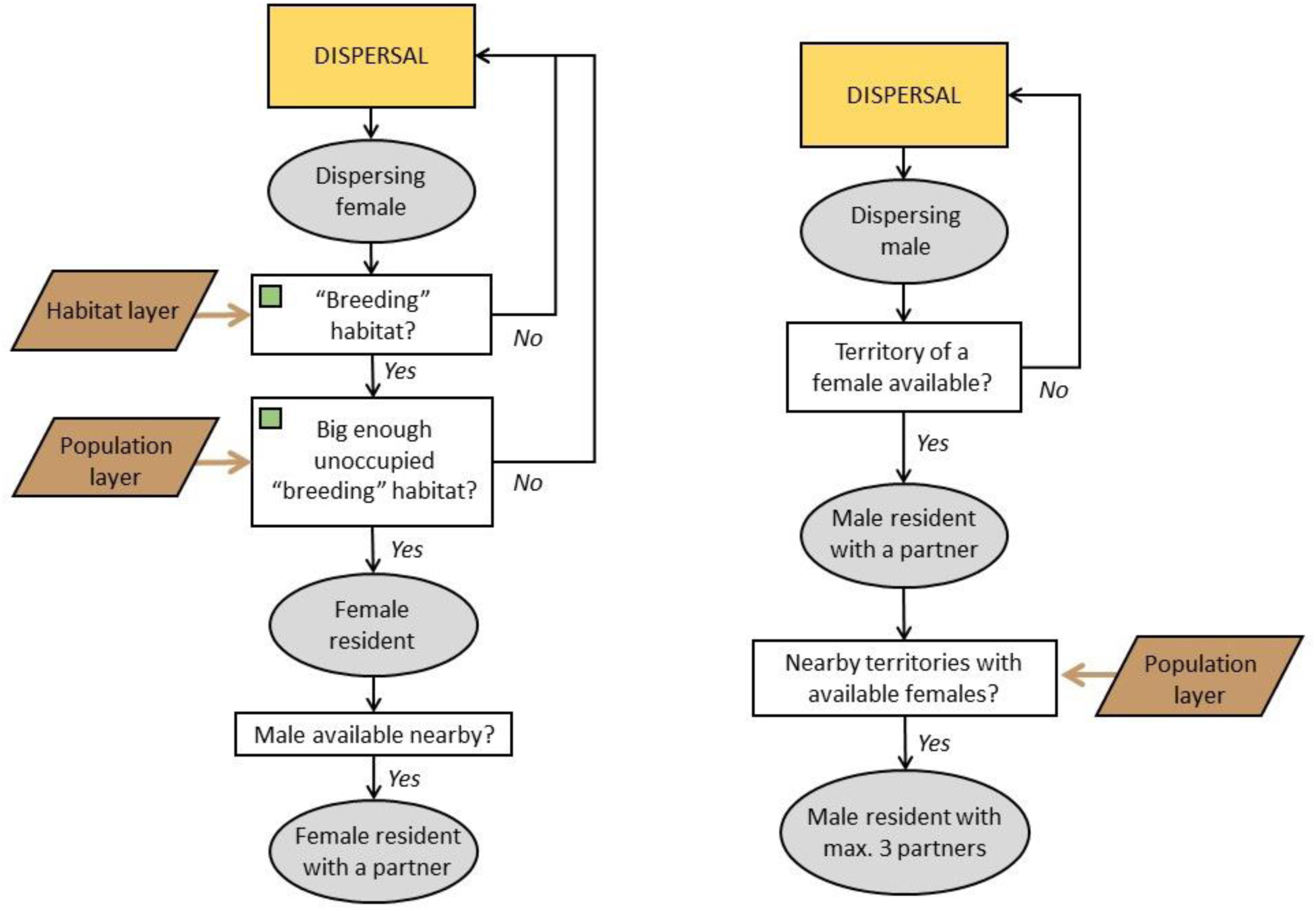
Diagram of the territory establishment in the lynx SE-IBM separated by sex. The successive grey circles represent the successive status of the individual. Females are constrained by their surrounding habitat defined by the “habitat layer” (brown box). Males are constrained by female presence. The territory sizes females need to establish in “breeding” habitat and the maximum distance to which males can look for available female residents are constrained by the population in which the individuals are located (“population layer”; brown box).

Lynx are solitary carnivores (i.e., each resident holds its own territory), except during reproduction (i.e., mating between male and female) and when females are raising their kittens (Breitenmoser et al., 1993; Stahl & Vandel, 1998). Sexually mature female residents reproduce if a sexually mature resident male occurs on their territory. Females have 0.5 chance to be sexually mature at one year old, and they are all sexually mature at two years old. Males have 0.5 chance to be sexually mature at two years old, all being sexually mature at three years old (Kvam, 1991; Kramer-Schadt, Revilla, & Wiegand, 2005; Breitenmoser-Würsten, Vandel, et al., 2007). In the wild, lynx litter size averages two kittens and can be up to four (Breitenmoser-Würsten et al., 2001; López-Bao et al., 2019). However, around 50% of lynx kittens die before reaching the age of dispersal (Breitenmoser-Würsten, Vandel, et al., 2007). We simulated that resident females up to 11 years old (“young” female in the SE-IBM model) produce one or two kittens, with a probability of 0.5 for each litter size, that will survive until becoming dispersers the following year (Henriksen et al., 2005; Kramer-Schadt, Revilla, & Wiegand, 2005). Senescence reduces litter size, therefore “old” females in the model (12 years old and older) produce zero or one kitten with an equal probability of 0.5 (Henriksen et al., 2005; Kramer-Schadt, Revilla, & Wiegand, 2005). We defined that residents die from a fixed annual mortality (i.e., baseline mortality, Table B.1) which does not include the mortality due to vehicle collisions (i.e., spatial mortality added separately with the collision model) and illegal killing (i.e., unavailable estimates). We did not define an increase of mortality due to senescence but we set an age maximum of 20 years (Stahl & Vandel, 1998; Breitenmoser-Würsten, Vandel, et al., 2007; von Arx et al., 2017). Residents can also die from a spatial annual mortality due to vehicle collisions inside their territory. This spatial mortality is specific to each resident and corresponds to mean collision probability inside their territory. A correction factor transforms the collision probability from the “collision layer” into a yearly collision mortality.

At the end of the year, all individuals (i.e., individuals which were disperser or resident at the beginning of the year) age by one year. Individuals still dispersing will do again this same loop and the new residents will do the annual loop, as long as the simulation lasts or until they die.

The model is coded using R 4.1.0 (R Core Team, 2014). We used the package NetLogoR (Bauduin, McIntire, & Chubaty, 2019) to facilitate the IBM structure implementation in R language and the package SpaDES (Chubaty & McIntire, 2018) to schedule SE-IBM rules with different time units. We also used the packages data.table (Dowle & Srinivasan, 2019), randomcoloR (Ammar, 2019), raster (Hijmans & Van Etten, 2018), and testthat (Wickham, RStudiom, & R Core Team, 2019).

### Appendix B Complete description of the lynx spatially-explicit individual-based model following the ODD protocol (Overview, Design concepts, and Details) developed by Grimm et al. (2006, 2010)

#### Overview

##### Purpose

The model simulates lynx population dynamics and dispersal, accounting for the impact of the road network via the risk of vehicle collisions and of the land cover to represent the lynx habitat preferences (Kramer-Schadt, Revilla, & Wiegand, 2005). Layers (i.e., maps) of collision probabilities, lynx habitats and lynx populations (Appendix A) are combined with SE-IBM rules simulating lynx dispersal (Kramer-Schadt et al., 2004, 2007) and demography (Kramer-Schadt, Revilla, & Wiegand, 2005; Kramer-Schadt et al., 2011).

##### Entities, state variables, and scales

The mobile entities of the model represent lynx individuals. Each simulated individual holds several characteristics:

- *id*: each lynx is unique and has a unique numerical identity (*id*);
- *heading*: direction of the lynx in degrees from 0 to 360 (0 is heading North) to one of its eight neighboring cells (i.e., 0°, 45°, 90°, 135°, 180°, 225°, 270°, 315°);
- *location*: coordinates of the lynx current position;
- *previous location*: lynx hold in memory the coordinates of their previous position;
- *population*: the population area where the individuals are born among Vosges-Palatinate, Black Forest, Jura or Alps;
- *sex*: male or female;
- *age*: numerical value to represent the lynx age. This number represents the age as for humans with zero for the first year of life, one for the second year of life, etc.;
- *status*: resident (i.e., established with a territory) or disperser (i.e., without a territory);
- *steps*: number of steps dispersing lynx have to do during the current day;
- *last dispersing location*: dispersing lynx hold in memory the coordinates of the last cell of habitat type “breeding” or “dispersal” they visited;
- *number matrix*: how many consecutive steps dispersing lynx move in “matrix” habitat;
- *male id*: for resident female only, the *id* of their male associated if they have one;
- *number females*: for resident male only, the number of resident females associated they have. It cannot be more than 3;
- *road mortality territory*: mean collision probability inside its territory;

Simulated lynx progress on a gridded study area of 1 km² resolution encompassing the four populations of interest: Vosges-Palatinian, Black Forest, Jura and Alpine, over Germany, France and Switzerland (main text, Fig. 1). The gridded study area resolution corresponds to the lynx perceptual range (Haller & Breitenmoser, 1986) and original resolution for the lynx individual-based models (Kramer-Schadt et al., 2004, 2011; Kramer-Schadt, Revilla, & Wiegand, 2005).

The gridded study area holds four different variables:

- *collision probability* (from the “collision layer”): cells crossed by roads have an estimated probability of fatal collision with lynx, a value between zero and one. The collision probability is equal to 0 for cells with no road in it;
- *habitat type* (from the “habitat layer”): each cell is of one of the following habitat types: “breeding”, “dispersal”, “matrix” or “barrier”;
- *individual territory*: this variable holds the *id* of the female residents for the cells included in their territories.

##### Process overview and scheduling

At the beginning of each simulated year, lynx individuals are differentiated by their resident or disperser *status* (Fig. A.4). First, dispersing lynx move on the gridded study area (Fig. A.5) on a daily time step. A number of *steps* to move per day is generated for each dispersing lynx at the beginning of each day. All dispersers move simultaneously, one step (i.e., cell) at the time, following a correlated habitat-dependent movement influenced by the *habitat type*. After each step, dispersing lynx may suffer from a spatial mortality given by the *collision probability* at their new location divided by a correction factor to transform the value into a per step collision mortality. Then, the surviving ones search for a place where establish their territory (Fig. A.6). Dispersing females are constrained by the *habitat type* in their surroundings to establish a new territory while dispersing males are constrained by the presence of available resident females for reproduction (Breitenmoser-Würsten, Zimmermann, et al., 2007b). Dispersing lynx that do not establish a territory continue to disperse during their set number of *steps* per day, and every day during the year until establish one. At the end of each day, lynx that dispersed suffer from a fixed daily baseline dispersal mortality. On the other hand, residents progress on a yearly time step. After the disperser movements, resident lynx may reproduce and kittens are born. Then, residents may die from a fixed annual baseline mortality and from a spatial annual mortality, the latter being defined by the mean *collision probability* inside their territories multiplied by a correction factor to transform the value into a yearly collision mortality. All these happen once in the year. At the end of a year, all lynx *age* are incremented by one and their *status* is updated if applicable.

Simulated lynx *id*, *population*, and *sex* variables do not change during a simulation. Their *heading*, *location*, *previous location*, *last dispersing location*, *number matrix*, *male id*, *number females* and *road mortality territory* may change at every dispersal step. The *steps* variable is updated each day, and *age* and *status* are updated once a year. There is no modeling of the landscape during the simulation, therefore the *collision probability* and *habitat type* are constant during one simulation. However, the *territory id* may change at every step (i.e., each time there is a new territory created).

#### Design concepts

##### Basic principles

Our lynx SE-IBM is an assemblage of four components: three spatial layers and an existing individual-based model (Kramer-Schadt et al., 2004, 2007, 2011; Kramer-Schadt, Revilla, & Wiegand, 2005) parameterized with field data and inverse model fitting (main text, Fig. 2 and Appendix C). The *collision probability* layer and the *habitat type* layer are output maps from a collision model and a habitat model (Appendix A). The “population layer” was generated using presence data and estimated number of lynx (Appendix A).

##### Emergence

Simulated males and females follow the same rules, except for the territory establishment where females are constrained by the habitat and males by the presence of females (Breitenmoser-Würsten, Zimmermann, et al., 2007b). In an area with few individuals, females have many possibilities of empty land to establish. On the other hand, males will have to move further to find one of the few females available, inducing longer dispersal movement and higher chances of dying. However, in a very dense area, this can be reverse. Females will have to move further to find free land whereas males, if they are not too numerous, may move less as they may have many females around them to pair with. If there are many males, all females may be already taken, inducing again longer dispersal for the males.

To mate, residents are constrained by the presence of a partner on their territory and both individuals need to be in age of reproduction. The presence of a suitable partner depends on their past dispersal movement and territory search, and if they could establish a new territory or not. Reproduction success is therefore stochastic, as well as the number of kittens. All this will modify the number of new individuals in the population.

Residents and dispersers are subject to a fixed baseline and spatial mortality. Both mortalities are stochastic and the spatial one depends on the location of the individuals (i.e., *collision probability* at their location). The location of the individuals depends on their current or past dispersal movement that depends on the *habitat type*. All this will modify the number of individuals dying each year.

##### Adaptation

There is no adaptation in the SE-IBM (e.g., no density-dependent rules). All individuals follow the same behavioral rules dependent on their inner and environmental characteristics.

##### Objectives

Simulated lynx do not have adaptive traits and they do not have a final goal to reach. They take decisions at each time step based only their current characteristics and their environment to fulfill an immediate goal (e.g., where to move next).

##### Learning

Simulated lynx try to minimize their time spent in poor habitat quality. When they step into “matrix” habitat, they store the position they just left (i.e., their last position in good habitat so either in “breeding” or “dispersing” habitat) and start counting how many consecutive steps into “matrix” they do. At the 10^th^ step, if the chosen cell is still located into “matrix”, simulated lynx will use their memory to find and move back to their last location of good habitat.

##### Prediction

Simulated lynx cannot predict the future nor have a global view of the study area. Both residents and dispersers sense the current state of their environment and population in their immediate surroundings and react to this information only. For example, dispersing lynx move, making decisions based only on their immediate surroundings. They cannot anticipate their path nor have a global view of their environment to make advantageous decisions for the long term (e.g., they cannot find a better territory and/or more quickly).

##### Sensing

Simulated lynx can sense the *habitat types* and the *individual territories* of their surroundings. Dispersers establishing a territory and trying to find a partner can sense other individuals: it knows who hold nearby territories and the characteristics of these individuals, such as their *sex* and the availability to pair with.

##### Interaction

There is a tacit competition between dispersing individuals during the territory search. Dispersers evaluate their surroundings before establishing one at the time (in a random order each time). The first female in the list looking for a territory has therefore more chance to find empty space to establish than the last female. Similarly, the first male in the list looking for an available female to join has more chance to find a partner than the last male.

Residents also interact with each other for reproduction. If a male and a female resident in age of reproduction are on the same territory, they may reproduce and produce new individuals. However, this interaction does not affect any of the individuals (i.e., a reproduction event does not change the characteristics of the concerned individuals).

##### Stochasticity

The model involves several stochastic processes. Regarding demography, both reproduction (i.e., the reproduction event itself and the number of kittens) and mortality (both fixed baseline and spatial for both residents and dispersers) are defined with probabilities and therefore outcomes vary between individuals. During dispersal, the choice of the *habitat type* to select when there is a mix of “breeding/dispersal” and “matrix” habitats in the lynx surroundings depends on the probability to step into “matrix”. Then, there is also stochasticity in the choice of the next location within the selected *habitat type*, as the choice to follow a correlated movement or not are defined by probabilities, as well as the choice of the next location among those available if the movement is not correlated. Finally, the number of steps dispersing simulated lynx must move during a day is sampled from a probability distribution and so, the outcomes are not the same for all individuals all the time.

##### Collectives

When a resident male finds one or several available resident females, he may pair and reproduce with them. Every year, the pairs persist unless one of the lynx dies and therefore frees the other one to find another partner.

Individuals belong to populations. The two rules different between populations are the size of the territory females have to reach and the maximum size of territory males can hold (i.e., how far can they look for females). Other than that, no rule is defined at the population level and the population affiliation does not affect the individual progress. Individual progress according to their own characteristics.

##### Observation

The state of all alive simulated individuals (i.e., their location and characteristics) and of the *individual territory* map are available at the end of each yearly time step. All disperser movements are also recorded on a single map over the whole simulation time. Events changing the demography are saved each year: which individuals died and how (i.e., from the fixed baseline mortality or from collision), which individuals reproduce, who are the kittens, and which individuals became residents.

#### Details

##### Initialization

At the beginning of the simulation, a grid covering the whole study area (main text, Fig. 1) with a resolution of 1 km^2^ is created. The values from the “habitat layer” and the “collision layer” are transferred to the gridded study area variables *habitat type* and *collision probability*. The variable *individual territory* is set to missing value (NA) everywhere. *Habitat type* on the borders of the grid are set to “barrier” so that individuals “bounce” on the borders. We assumed that these individuals would have had a hard time to go beyond the borders of the study area as we encompassed the main populations and patches of good habitats and therefore would be unlikely to leave the study area.

Using the “population layer”, a lynx population is created with unique *id* for each individual, *location*, *population*, as well with a *sex* and *age* (when known from field data) for some individuals. A simulated individual cannot start in “matrix” or “barrier” habitats. Therefore, if some individuals’ *location* are located in these habitats, they are relocated to the closest cell of “breeding” or “dispersal” habitat. At the start of the simulation, individuals cannot have a *previous location* or a *last dispersing location*, so we used their current *location* for these two variables. For individuals with unknown *sex* or *age* (no field data), a random *sex* (male or female with ratio 1:1) and *age* (between 2 and 15) is randomly given. A random *heading* (i.e., initial direction) is given to each individual, between 0° and 360°. At the start, all individuals have a disperser *status*, 0 *steps* to do, 0 steps done in “matrix” (*number matrix*) and 0 for *road mortality territory*. As all individuals are dispersers, no couples are made so *male id* for the females are set to NA and the *number females* for males are set to 0 (Table B.1).

The parameter values used in the SE-IBM to represent the dynamics and dispersal of the Eurasian lynx in our study area (main text, Fig. 1) are provided in Table B.1.

**Table B.1:** Parameter values used in the SE-IBM.

| Parameter | Explanation | Value | Reference |
| --- | --- | --- | --- |
| endTime | Number of years to simulate the lynx populations. | 50 | By default |
| pRepro | Probability of mating for a resident female which has a resident male associated on her territory when they are both in age of reproduction. | 0.95 | Model calibration (see Appendix C) |
| nKittyYoungF | Number of kittens that survive until the age of dispersal when the female is considered “young” (maxAgeYoungF parameter). | 1 or 2<br>(with equal probability for each value) | (Breitenmoser-Würsten et al., 2001; Henriksen et al., 2005; Kramer-Schadt, Revilla, & Wiegand, 2005; Breitenmoser-Würsten, Vandel, et al., 2007; López-Bao et al., 2019) |
| nKittyOldF | Number of kittens that survive until the age of dispersal when the female is considered “old” (maxAgeYoungF parameter). | 0 or 1<br>(with equal probability for each value) | (Breitenmoser-Würsten et al., 2001; Henriksen et al., 2005; Kramer-Schadt, Revilla, & |
|  |  |  | Wiegand, 2005;<br>Breitenmoser-Würsten,<br>Vandel, et al.,<br>2007) |
| maxAgeYoungF | Age maximum for a female to be considered “young”. | 11 | (Henriksen et al., 2005) |
| minAgeReproF | Age minimum at which all resident females can reproduce (i.e., are sexually mature). One year younger, only half of the females are mature. | 2 (3<sup>rd</sup> year of life) | (Kvam, 1991;<br>Henriksen et al., 2005;<br>Breitenmoser-Würsten,<br>Vandel, et al., 2007) |
| minAgeReproM | Age minimum at which all resident males can reproduce (i.e., are sexually mature). One year younger, only half of the males are mature. | 3 (4<sup>th</sup> year of life) | (Kvam, 1991) |
| pMortResAlps | Fixed annual baseline probability of mortality for resident lynx (without the risk of collisions included) of the Alpine population. | 0.15 | Model calibration (see Appendix C) |
| pMortResJura | Fixed annual baseline probability of mortality for resident lynx (without the risk of collisions included) of the Jura population. | 0.15 | Model calibration (see Appendix C) |
| pMortResVosgesPalatinate | Fixed annual baseline probability of mortality for resident lynx (without the risk of collisions included) of the Vosges-Palatinian population. | 0.15 | Same value as pMortResJura, see model calibration (Appendix C) |
| pMortResBlackForest | Fixed annual baseline probability of mortality for resident lynx (without the risk of collisions included) of the Black Forest population. | 0.15 | Same value as pMortResJura, see model calibration (Appendix C) |
| corrFactorRes | Correction factor for road mortality risk for residents. This factor multiplies the *road mortality territory* value to transform it into a yearly collision mortality. | 3.9 | Model calibration (see Appendix C) |
| corrFactorDisp | Correction factor for road mortality risk for dispersers. This | 170 | Model calibration (see |
|  | factor divides the collision probability of from the “collision layer” to transform the value into a per step collision mortality. |  | Appendix C) |
| ageMax | Age maximum a lynx can reach.<br>When reaching this age, lynx die from natural causes. | 20 | (von Arx et al., 2017) |
| xPs | Exponent value of the power function to define the daily step distribution. This distribution is used to sample a number of steps per day to move for dispersing lynx. | 11 | (Kramer-Schadt et al., 2004) |
| sMaxPs | Maximum number of steps dispersing lynx can do in one day. | 45 | (Kramer-Schadt et al., 2004) |
| pMat | Probability of stepping into “matrix” cells. This probability is multiplied by the number of “matrix” cells available for a dispersing lynx to define its probability to choose this habitat type for its next location. | 0.03 | (Kramer-Schadt et al., 2004) |
| pCorr | Movement correlation probability | 0.5 | (Kramer-Schadt |
|  | for dispersal movement. |  | et al., 2004) |
| pMortDispAlps | Fixed daily baseline probability of mortality for dispersing lynx (without the risk of collisions included) of the Alpine population. | 0.0007 | Model calibration (see Appendix C) |
| pMortDispJura | Fixed daily baseline probability of mortality for dispersing lynx (without the risk of collisions included) of the Jura population. | 0.0009 | Model calibration (see Appendix C) |
| pMortDispVosgesPalatinate | Fixed daily baseline probability of mortality for dispersing lynx (without the risk of collisions included) of the Vosges-Palatinian population. | 0.0009 | Same value as pMortDispJura, see model calibration (Appendix C) |
| pMortDispBlackForest | Fixed daily baseline probability of mortality for dispersing lynx (without the risk of collisions included) of the Black Forest population. | 0.0009 | Same value as pMortDispJura, see model calibration (Appendix C) |
| nMatMax | Maximum number of consecutive steps a dispersing lynx can do into “matrix” habitat. After this number of steps, disperser can use its memory to find and move in the last good habitat visited (“breeding” or “dispersal” habitat). | 9 | (Kramer-Schadt et al., 2004) |
| coreTerrSizeFAlps | Territory size (in km<sup>2</sup>) females located in the Alpine population need to reach. | 76 | (Breitenmoser-Würsten et al., 2001) |
| coreTerrSizeFJura | Territory size (in km<sup>2</sup>) females located in the Jura population need to reach. | 119 | (Breitenmoser-Würsten, Zimmermann, et al., 2007b) |
| coreTerrSizeFBF | Territory size (in km<sup>2</sup>) females located in the Black Forest population need to reach. | 119 | (Breitenmoser-Würsten, Zimmermann, et al., 2007b) |
| coreTerrSizeFVP | Territory size (in km<sup>2</sup>) females located in the Vosges-Palatinian population need to reach. | 119 | (Breitenmoser-Würsten, Zimmermann, et al., 2007b) |
| terrSizeMAlps | Maximum territory size (in km<sup>2</sup>) males located in the Alpine population can reach. | 137 | (Breitenmoser-Würsten et al., 2001) |
| terrSizeMJura | Maximum territory size (in km<sup>2</sup>)<br>males located in the Jura<br>population can reach. | 226 | (Breitenmoser-Würsten,<br>Zimmermann, et<br>al., 2007b) |
| terrSizeMBF | Maximum territory size (in km<sup>2</sup>)<br>males located in the Black Forest<br>population can reach. | 226 | (Breitenmoser-Würsten,<br>Zimmermann, et<br>al., 2007b) |
| terrSizeMVP | Maximum territory size (in km<sup>2</sup>)<br>males located in the Vosges-Palatinian population can reach. | 226 | (Breitenmoser-Würsten,<br>Zimmermann, et<br>al., 2007b) |
| nFem | Maximum number of resident<br>females a resident male can pair<br>with and potentially reproduce<br>with. | 3 | (Kramer-Schadt,<br>Revilla, &<br>Wiegand, 2005) |

##### Input data

See Appendix A.

##### Submodels

Dispersal: A number of steps to do during the day is sampled every day from the distribution defined by Kramer-Schadt et al. (2004), one per dispersing individual. Dispersers move one cell per step, into one of their eight neighboring cell or their current cell (i.e., so a total of nine potential cells for their next location). First, there is a selection of the habitat type each disperser chooses for its next location. Among those nine cells, all cells of “barrier” type are removed from the potential choices. As individuals cannot be on “barrier” habitat, it cannot happen that all nine cells are “barrier”. For a dispersing individual, if all his cells are of “breeding” or “dispersal” type, this type (“breeding/dispersal” without distinction between the two) is selected. If all his cells are of “matrix” (i.e., habitat not favorable for lynx), this type is selected. If the cells are a mix of “breeding/dispersal” and “matrix”, there is a Bernoulli trial of mean 0.03 (Table B.1) times the number of “matrix” cells among the nine to select the “matrix” habitat type.

Second, there is the choice of the cell for the next location, among all cells of the selected habitat type. If there is only one cell of the selected type, the disperser goes on this one (for example, if the selected cell type is “breeding/dispersal” and there is only one cell of “breeding/dispersal” habitat, all other cells being “matrix” or “barrier”, therefore the only cell of “breeding/dispersal” is selected). Otherwise, there is a choice either random or governed by a correlation movement to select the cell. The first step of the day is never correlated, otherwise there is a Bernoulli trial of mean 0.5 (Table B.1) to determine if the movement is correlated for each individual for each step. If there is no correlation of the movement, the choice for the cell is done with an equal probability for all the cells of the selected type. If the movement is correlated, the rotating angle between the individual’s heading and each cell of the selected type is calculated. There is a preference for the cell minimizing this rotating angle. If the cell that the individual is on is of the selected habitat type (i.e., this cell can be selected for the individual’s next step which will make the lynx stay at its current location), the preference for this cell is equal to the preference for the cells inducing a rotation of +/− 90 degrees, to favor a forward movement. The selected cell is the one with the highest preference (i.e., smallest rotation movement).

Third, if the selected cell is of “matrix” type, the individuals check how many consecutive steps they have done in “matrix” habitat (*number matrix*). If a disperser has already step nine consecutive times (Table B.1) in “matrix” and this 10^th^ consecutive step is in “matrix” too, it used its memory and select, for its next location, the last visited cell of good habitat (“breeding” or “dispersal”), stored in his memory (*last dispersing location*) (Kramer-Schadt et al., 2004).

Finally, all dispersers rotate towards their selected cell and move to the center of it. They update their *previous location* with the coordinates of the cell they were before moving. If their new location is of type “breeding” or “dispersal”, this location is stored in their memory for the *last dispersing location*, and their *number matrix* is reset to zero. If their new location is of type “matrix”, their *number matrix* increments of one.

Now, disperser may be also subject to spatial mortality. There is a Bernoulli trial for each disperser with mean equal to the *collision probability* value of their new location divided by a correction factor (Table B.1). Dispersers that do not die from spatial mortality search for a new home range to establish (*searchTerritory* sub-model).

The dispersal movement is done until all dispersing individuals have reached their number of steps to do during the day, or they have established a territory and are now resident, or they died from spatial mortality. At the end of the day, all individuals that were dispersers at the beginning of the year (even the ones that established a territory during the day or previous day) may die from a fixed daily baseline mortality using a Bernoulli trial for each individual. A daily spatial mortality is also applied to individuals which dispersed during the year but already found a territory during previous days. As movements inside territories are not simulated but they were not considered as resident at the beginning of the year, a per step or an annual spatial mortality cannot be applied. Their daily spatial mortality is then calculated as their spatial mortality with their resident status divided by 365. If some of the individuals dying established a territory during this day, there is an update of their characteristics similarly as the death of resident individuals (*mortality* submodel).

At the beginning of the simulation, all initial individuals were created as dispersers to let simulated individuals establish their initial territories. Individuals needs some time to do so, therefore we did not apply any mortality (daily baseline nor spatial) during the first year of simulation to allow dispersers to establish their territories.

SearchTerritory: Dispersing individuals arriving at a new location search for a territory to establish, one at the time, and females first (i.e., male establishment depending on female territories). One at the time, in a random order each time, females evaluate their surroundings to establish their territory. Females needs to be on a cell of “breeding” type that is not already included in any territory to start creating their own. Then, they need to have enough empty contiguous cells of “breeding” habitat (search in 8-cell directions) to create a big enough territory (Table B.1). When enough empty contiguous cells of “breeding” habitat are present, females establish their territory and becomes resident. Otherwise, they stay disperser and keep dispersing. After a female establishes her territory, all the cells of this territory become occupied and are not available anymore for the other females. The female *id* is given to the *individual territory* study area variable for all cells in her territory. The mean *collision probability* of the territory is calculated and given to the female for her *road mortality territory*. Then, the female checks if there is a resident male around. If there is a male not further than the radius of the maximum territory size for the male (Table B.1) and he has less than three females associated, she pairs with him. Then, she obtains the male *id* for her *male id* and the *number females* of the male increments of one.

After all dispersing females tried to establish, dispersing males try to disperse of their own, one at the time in a random order each time. Each male checks if the cell they are currently on is located inside the territory of an available female (i.e., without male associated yet). If there is one, the male pairs with this female and becomes resident. His territory is the same one as the female and its *number females* is set to one. The female obtains the male *id* for her *male id* variables. Then, the male also looks if there are other nearby available females to pair with, as far as the radius of its maximum territory size (Table B.1). He can pair with up to three available females. The male territory represents the union of all the female territory with which he paired with. His *road mortality territory* is the mean value of the *road mortality territory* of all the females he paired with.

Reproduction: A female resident in *age* of reproduction (one or two years old, see Table B.1) with a resident male associated, also in *age* of reproduction (two or three years old, see Table B.1), may reproduce with a Bernoulli trial (Table B.1) and have offspring. If the couple reproduces, the female can produce one or two kittens with equal probability (Table B.1) if she is “young” (11 years old or younger; Table B.1) or zero or one kitten with equal probability if she is “old” (12 years old or older; Table B.1). Kittens obtain a unique *id*. The *sex* of the kittens is randomly chosen between male and female with ratio 1:1 (Table B.1). Their *location*, *heading* and *last dispersing location* are the ones from their mother. Their *previous location* is NA. Their *population* is not the one of their parents but the one of the territory where they are born. Their *age* is 0, their *status* is resident (i.e., they stay with their mother and cannot disperse at this age), their *steps* and *number matrix* are zero, their *male id* is NA, and their *number females* and *road mortality territory* are zero.

Mortality: Resident individuals may die from a fixed baseline annual mortality (Table B.1) and a spatial mortality using a Bernoulli trial. We did not simulate an effect of senescence but we set an age maximum (20 years; Stahl & Vandel, 1998; Breitenmoser-Würsten, Vandel, et al., 2007; von Arx et al., 2017) and all individuals reaching this age die from a non-collision source (i.e., old lynx mortality is not included in the collision mortality). Resident individuals may also die from spatial mortality, the mortality rate for female residents is the mean *collision probability* of the cells inside of their territory (*road mortality territory*) multiplied by a correction factor (Table B.1). For male residents, spatial mortality rate is the mean *road mortality territory* of all his paired females multiplied by the same correction factor (Table B.1).

When a resident female dies, her territory disappears (i.e., her *id* is removed from the *individual territory* study area variable) and the local male resident loses one female. If this male did not have other females, he becomes disperser again to look for new available female resident as male home ranges are adjusted to mate availability (Breitenmoser-Würsten, Zimmermann, et al., 2007b). If the dead female had kittens during the year, they all die too. When a male resident dies, its paired females become available to reproduce with other males.

Demography: All individuals’ *age* increment of one. The *status* of the offspring of the year changes from resident to disperser.

### Appendix C Calibration of the parameters, sensitivity analysis and model validation

#### Calibration of the parameters

We calibrated the lynx SE-IBM using a pattern-oriented modeling (Grimm & Railsback, 2012; Gallagher et al., 2021) in a two-phase process. To better adapt the model to our study area and simulated lynx populations, we (re)estimated the following 7 parameters:

- pMortResAlps / pMortDispAlps / pMortResJura / pMortDispJura: Fixed annual baseline mortalities (i.e., probabilities) applied to resident / dispersing lynx, without the risk of collisions included, for the Alpine and the Jura populations. These parameters do not exactly represent the mortality rates for lynx when omitting collision mortality observed in the field as the mortality due to old age in our model is additive to these estimated baseline probabilities. For the mortality probabilities for the Vosges-Palatinian and the Black Forest populations, we used those of the Jura population and they were not calibrated *per se*;
- corrFactorRes: Correction factor for road mortality risk for residents. This factor multiplies the road mortality territory to transform the value into a yearly collision mortality;
- corrFactorDisp: Correction factor for road mortality risk for dispersers. This factor divides the collision probability from the “collision layer” to transform the value into a per step collision mortality;
- pRepro: Probability of mating for a resident female which has a resident male associated on the same territory, when they are both in age of reproduction. This parameter is different than the probability of reproducing observed in the field, detected by the presence of kitten, as, in our model, old female can mate but produce 0 young because of her age.

For the first phase of calibration, we sampled one value for each parameter to calibrate among wide ranges of values (Table C.1). We defined 50 calibration scenarios by sampling a value within the range for each parameter for each scenario. We ran 15 replicates of each calibration scenario. We ran the lynx SE-IBM forecasting the populations over 50 years and we used a different initial population (Appendix A) at each replicate within each scenario to avoid bias due to initial locations of simulated individuals. We did not simulate any mortality during the first year of simulation to allow dispersing individuals to find a territory, if they can, without dying while doing so. From simulation outputs, we computed 13 patterns for each scenario, taking the mean value over the 15 replicates. Patterns were population emergent properties from simulations that we could compare to reference values. We defined a burn-in phase of 2 years after the start of the simulation to let the population settle down so that only outputs from the 3^rd^ year of simulation were used to compute patterns. Reference values for the patterns were extracted from literature or computed from field data. In order to calibrate the model, we looked for the calibration scenarios (i.e., and their associated parameter values) which minimized the differences between the pattern values extracted from their simulation outputs and the pattern values from external sources. The patterns we tried to match with their values and 95% confidence intervals from literature or data were:

- rAllRes_Alps = 0.24 [0.08;0.57] / rAllDisp_Alps = 0.24 [0.15;0.38]: Annual mortality rates observed for resident / dispersing individuals from all causes of mortality (i.e., including collisions) for the Alpine population (Premier et al., 2025).
- rCollRes_Alps = 0.02 [0;0.059] / rCollDisp_Alps = 0.077 [0;0.211]: Annual mortality rates observed for resident / dispersing individuals from collisions with vehicles for the Alpine population (Premier et al., 2025).
- nCollAlps = 2 [0;3]: Mean annual number of fatal collisions observed for the Alpine population (data for 2017-2021: OFB Réseau Loup-Lynx and KORA).
- rAllRes_Jura = 0.17 [0.1;0.29] / rAllDisp_Jura = 0.35 [0.16;0.65]: Annual mortality rates for resident / dispersing individuals from all causes of mortality (i.e., including collisions) observed for the Jura population (Premier et al., 2025).
- rCollRes_Jura = 0.034 [0;0.08] / rCollDisp_Jura = 0.067 [0;0.185]: Annual mortality rates for resident / dispersing individuals from collisions with vehicles observed for the Jura population (Premier et al., 2025).
- nCollJura = 6 [3;9]: Mean annual number of fatal collisions observed for the Jura population (data for 2017-2021; OFB, Réseau Loup-Lynx and KORA).
- rRepro = 0.81: Observed proportion of resident females reproducing (Breitenmoser-Würsten, Vandel, et al., 2007a; López-Bao et al., 2019).
- distDispAlps = 26 [15;36] / distDispJura = 63 [39;87]: Mean dispersal distance in kilometers, computed as a straight line, between the natal territory and the new established territory after dispersal observed for the Alpine and the Jura population (Zimmermann, Breitenmoser-Würsten, & Breitenmoser, 2005).

We removed from the calibration process the calibration of the parameter pRepro and its associated pattern rRepro as quick tests showed that a pRepro of 0.95 produced a stable mean of rRepro equal on average to 0.79 [0.78;0.80], independently from the other parameter values. Among the 50 calibration scenarios, none produced patterns for which the mean values computed from the simulation outputs fall into the 95% confidence intervals of the reference values for all patterns simultaneously. We therefore omitted the patterns about dispersal distances for this step (distDispAlps and distDispJura) as we chose to focus on the mortality process. Without these two patterns about dispersal distances, 2 out 50 scenarios produced patterns within the confidence intervals for the 10 remaining patterns (Table C.2).

For the second phase, we fined tuned the calibration of the 6 parameters to calibrate (i.e., still omitting rRepro). We explored a narrower range of values for each parameter, which included the parameter values sampled in the two scenarios that resulted in the best calibrations in the first phase (Table C.1). Similarly as in the first phase, we ran 50 new calibration scenarios (with 15 replicates each time) in the same conditions, and computed the same patterns. Again, none of the scenarios could satisfy the 13 patterns simultaneously. When removing the reproduction (i.e., not relevant) and the dispersal distances (i.e., too constraining), 25 scenarios produced patterns with their mean values falling within the confidence intervals of the reference values for the 10 remaining patterns. We finally selected, among those 25, the scenario which produced patterns with mean values the closest to the 13 pattern values simultaneously (Table C.2). The parameter values sampled for this scenario (Table C.1) were the ones selected and used to run the simulations to produce the results presented in the main text.

**Table C.1:** Value tested or selected for the parameters to be calibrated. “Phase 1” and “Phase 2” present the range of the values tested for each parameter for the two phases of the calibration. One value was randomly sampled within each range for each calibration scenario. “Phase 1. Best 1” and “Phase 1. Best 2” are the values from the best two scenarios of the first phase of calibration. “Selected” presents the selected values at the end of the phase 2 of the calibration process.

|  | pMortResAlps | pMortDispAlps | pMortResJura | pMortDispJura | corrFactorRes | corrFactorDisp |
| --- | --- | --- | --- | --- | --- | --- |
| Phase 1 | [0.05;0.25] | [0.0001;0.002] | [0.05;0.25] | [0.0001;0.002] | [1;6] | [50;300] |
| Phase 1. Best 1 | 0.15 | 0.0007 | 0.19 | 0.0008 | 3.5 | 175 |
| Phase 1.<br>Best 2 | 0.15 | 0.001 | 0.12 | 0.0007 | 3.5 | 175 |
| Phase 2 | [0.13;0.17] | [0.0005;0.0012] | [0.10;0.21] | [0.0005;0.001] | [3;4] | [125;225] |
| Selected | 0.15 | 0.0007 | 0.15 | 0.0009 | 3.9 | 170 |

**Table C.2:**
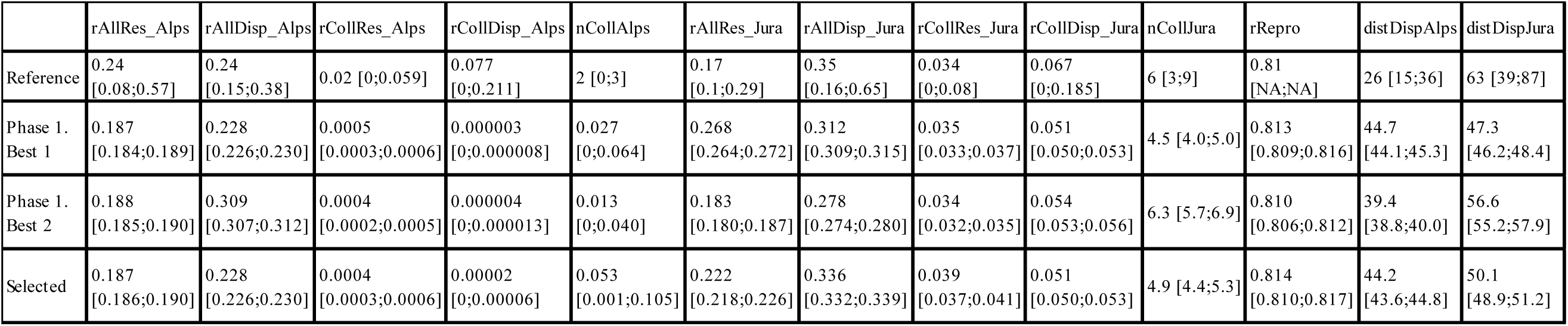
Patterns values. “Reference” are values (mean and 95% confidence intervals) extracted from the literature or field data. “Phase 1. Best 1” and “Phase 1. Best 2” are the pattern values (mean and 95% confidence intervals over the 15 replicates) computed with the simulation outputs from the scenarios which were the best ones from the first phase of the calibration process. “Selected” presents the pattern values (mean and 95% confidence intervals over the 15 replicates) computed with the simulation outputs from the best calibration scenario selected in the second phase of the calibration process.

#### Sensitivity analysis

The second phase of the calibration was used to perform the sensitivity analysis. Multiple values were tested for each parameter, values smaller and larger than the ones selected. We compared the percentage of variation of the parameter values used in the second phase of the calibration (Fig. C.1) with the percentage of variation on the main result of the simulation, the population growth rates (Fig. C.2).

**Figure C.1:**
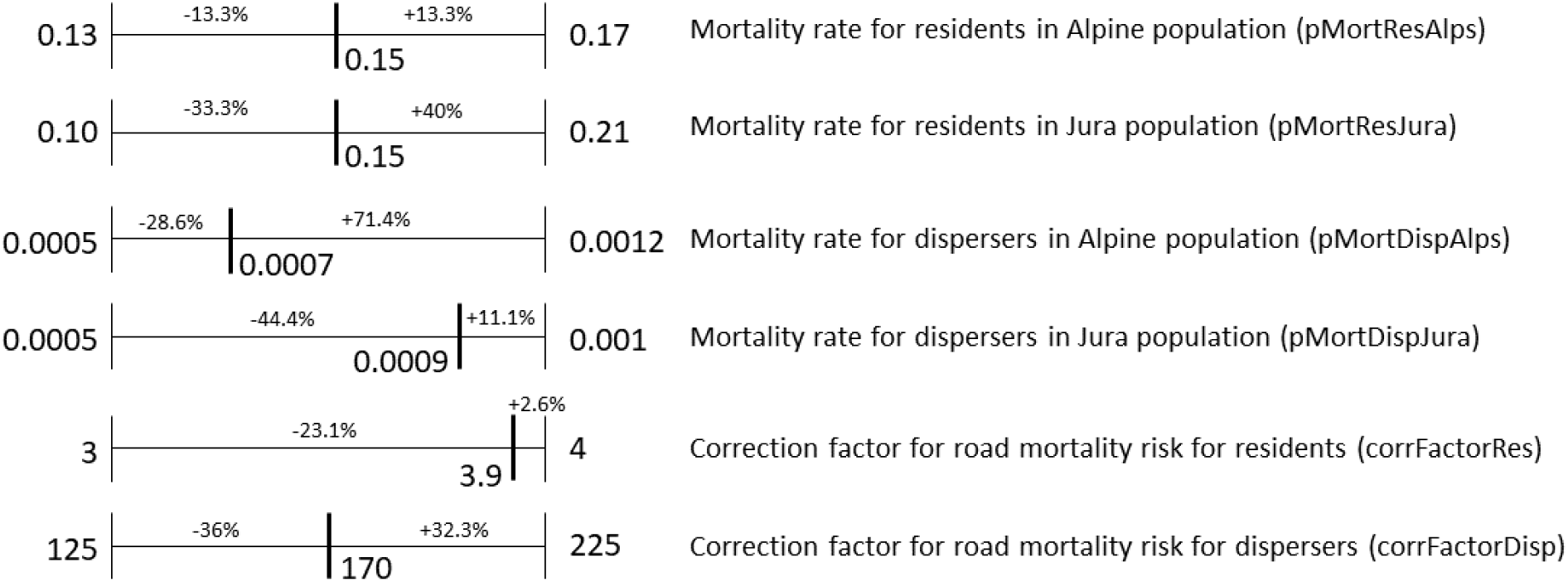
Range of values tested during the second phase of calibration for each calibrated parameter. The value indicated within the range of each parameter was the selected one with the best calibration scenario. Percentages noted are the difference between the selected value and the minimum and maximum values of the range for each parameter.

**Figure C.2:**
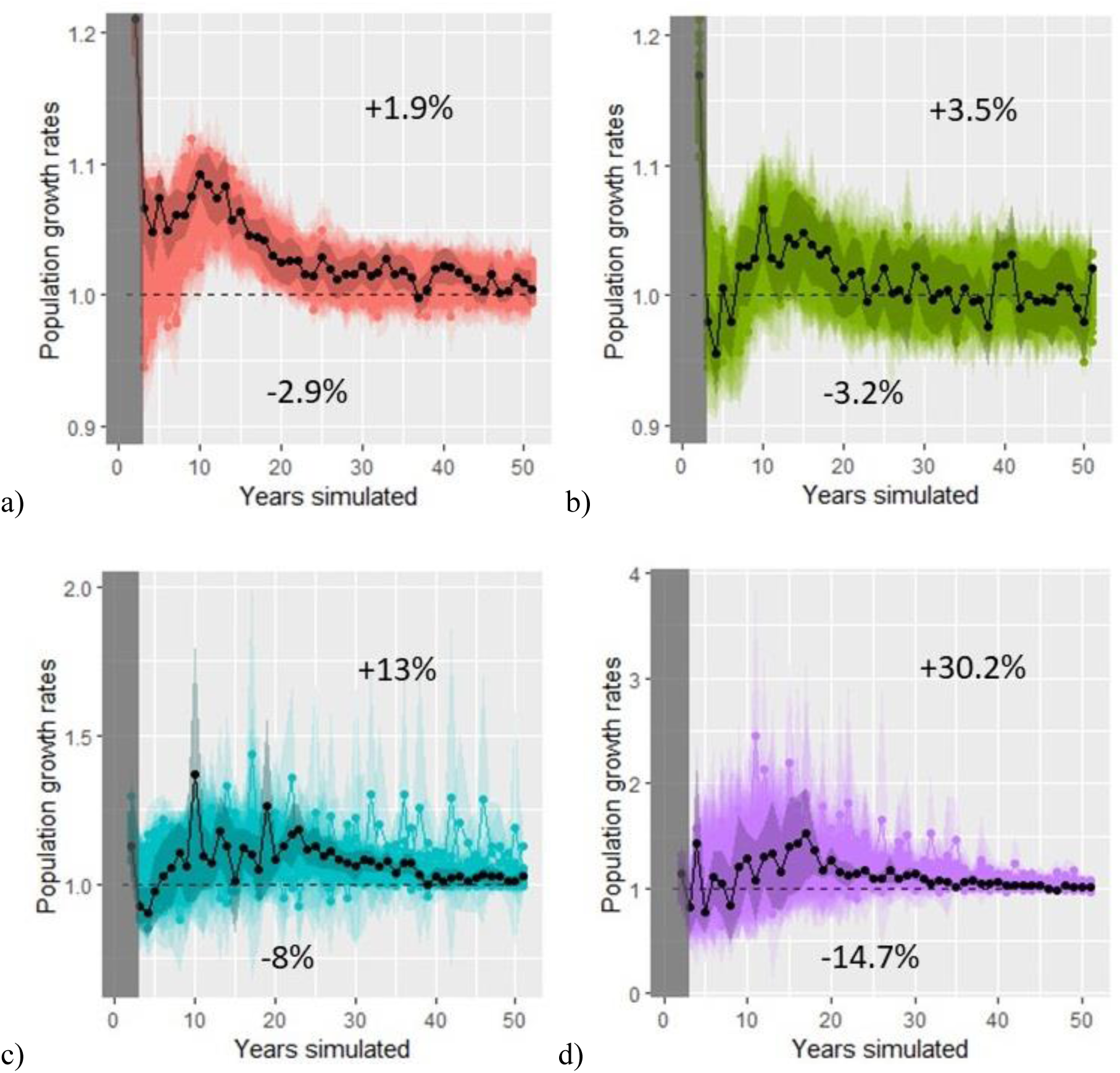
Annual rates of increase over the simulated years for each population (a) Alpine, b) Jura, c) Vosges-Palatinian, d) Black Forest). The grey area represents the 2-year burn-in phase at the beginning of the simulation. Overlaid colored points are mean over the 15 replicates and overlaid colored envelopes are 95% confidence intervals from the 15 replicates, for each of the 50 calibration scenarios from the second calibration phase. The black point and darker colored envelope represent the mean and 95 % confidence interval from the 15 replicates for the best calibration scenario selected during this second calibration phase. The dashed line represents a stabilization of the population (growth rate equals to 1). The percentage noted above the lines is the mean over all the years of the differences between the values for the selected scenario (means over the 15 replicates) and the maximum values of each year (maximum means calculated over the 15 replicates for the particular scenario this year). The percentage noted below is the same value but calculated with the minimum values each year.

The growth rates calculated for the Alpine and the Jura population were not sensitive to the different parameter estimates as the variations in parameter values were much larger than those of the result values. Parameters varied from −13.3% down to −44.4% and from +2.6% up to 40% in values (Fig. C.1), whereas the growth rates for these two populations varied on average from −2.9% up to +1.9% for the Alpine population and from −3.2% up to +3.5% for the Jura population (Fig. C.2). The variation in the growth rates for the Vosges-Palatinian population were a little larger (from −8% up to +13%) (Fig. C.2) but still smaller than most of the variations of the parameters (Fig. C.1). The growth rates for the Black Forest population on the other hand seemed sensitive to the model parameters with variations from −14.7% up to +30.2% (Fig. C.2). Results concerning this population were therefore less robust to interpretation. However, this population had very large confidence intervals regarding it growth rates compared to the other populations which made the confidence in the population trend already less precise for interpretation (main text, Fig. 3). The Vosges-Palatinian population and especially the Black Forest one had very few starting individuals (i.e., more prone to stochasticity) and data from the literature, making the model harder to calibrate regarding these populations and therefore less reliable to forecast their future. In comparison, the Alpine and Jura populations had more starting individuals and data from the literature to calibrate the model and therefore showed a stronger confidence and precision in the results.

#### Model validation

Among the 13 patterns used for calibration, the selected set of parameters (i.e., calibration scenario selected during the second phase) produced simulation results for which the pattern values of 11 of them fell into the 95% confidence intervals of the reference values (Table C.2). However, the confidence intervals for the pattern simulated values were extremely narrow (Tables C.2 and C.3) compared to those of the reference values. Therefore, model stochasticity did not seem very important. The reproduction rate reference value did not have a confidence interval available but the simulated value was very close and the reference value was inside the 95% confidence interval of the simulations. We completed this validation phase looking at the number of simulated collisions for the first few years with the concurrent field data. As the model was parameterized with 2017 data, we compared the simulated collisions for the Alpine and Jura populations for the first five years with the recorded collisions in these two populations for 2017 until 2021 (Table. C.3). From the calibration pattern results and this collision validation, we were very confident in the selected parameter values regarding the demographic parameters (i.e., mortality and reproduction rates), and especially for the Jura population.

The simulated collisions data matched less the field data for the Alpine, although the number of collisions were very low in this population (Table C.3). Also, the only discrepancy between the reference values and the simulated results for the calibration patterns (Table C.2) were for the dispersal distances for the Alpine population. The individuals seemed to have dispersed further in the simulations than computed from the field data. The model may need more attention on the interpretation for the Alpine population as the mortality may be slightly under-estimated and dispersal distances slightly over-estimated. Both could be linked as individuals could move further away as they were dying less doing so.

**Table C.3:** Number of vehicle collisions with adult or subadult lynx. Collisions with juveniles were not included in the recorded data as the SE-IBM did not simulate movement for the lynx in their first year of life and therefore cannot reproduce juvenile collisions. “Recorded” were real collisions recorded in the field. “Simulated” were collisions simulated with the 100 simulation replicates using the parameter values selected (mean and 95% confidence interval over the 100 replicates).

|  | **2017** | **2018** | **2019** | **2020** | **2021** |
| --- | --- | --- | --- | --- | --- |
| Number of collisions recorded for the Jura population | 6 | 10 | 4 | 5 | 5 |
| Number of collisions simulated for the Jura population | 5.7<br>[5.1;6.2] | 5.1<br>[4.6;5.6] | 5.5<br>[5.1;5.9] | 5.3<br>[4.8;5.8] | 4.8<br>[4.4;5.2] |
| Number of collisions recorded for the Alpine population | 2 | 0 | 3 | 1 | 3 |
| Number of collisions simulated for the Jura population | 0.01<br>[0;0.03] | 0.03<br>[0;0.06] | 0.03<br>[0;0.06] | 0.01<br>[0;0.03] | 0.04<br>[0;0.08] |

### Appendix D Model validation using telemetry data

We performed a qualitative validation of the lynx model predictions using tracks from collared female residents. The map below (Fig. D.1) shows the simulated territory occupancy with darker red colors representing areas most often occupied by female territories across simulation replicates and blue the least occupied (main text, Fig. 5). We overlaid GPS and VHF paths from collared female residents on the map. Data in the Vosges-Palatinate consist in two females in the Palatinate Forest (Germany) and one in the Vosges Mountains (France) (locations recorded in 2017-2018; Scheid et al., 2021). Data in the Jura represent four females in France and 13 in Switzerland (locations recorded in 1988-1998; Breitenmoser-Würsten, Zimmermann, et al., 2007b). Data in the Alps (Switzerland) represent 16 females (locations recorded in 1997-2000; Breitenmoser-Würsten et al., 2001). We show the full study area and four zoomed areas on the telemetry locations. Overall, the realized movements and territory emergence inferred from field data were coherent with the frequencies across simulation replicates of the territories defined by simulated lynx. Very few telemetry locations fall in white and grey cells (i.e., non-breeding habitat and breeding habitat cells never selected as part of a female territory across the 100 simulation replicates respectively) and most of the recorded tracks were located in cells with a high frequency of territory occupancy across the simulation replicates.

**Figure D.1:**
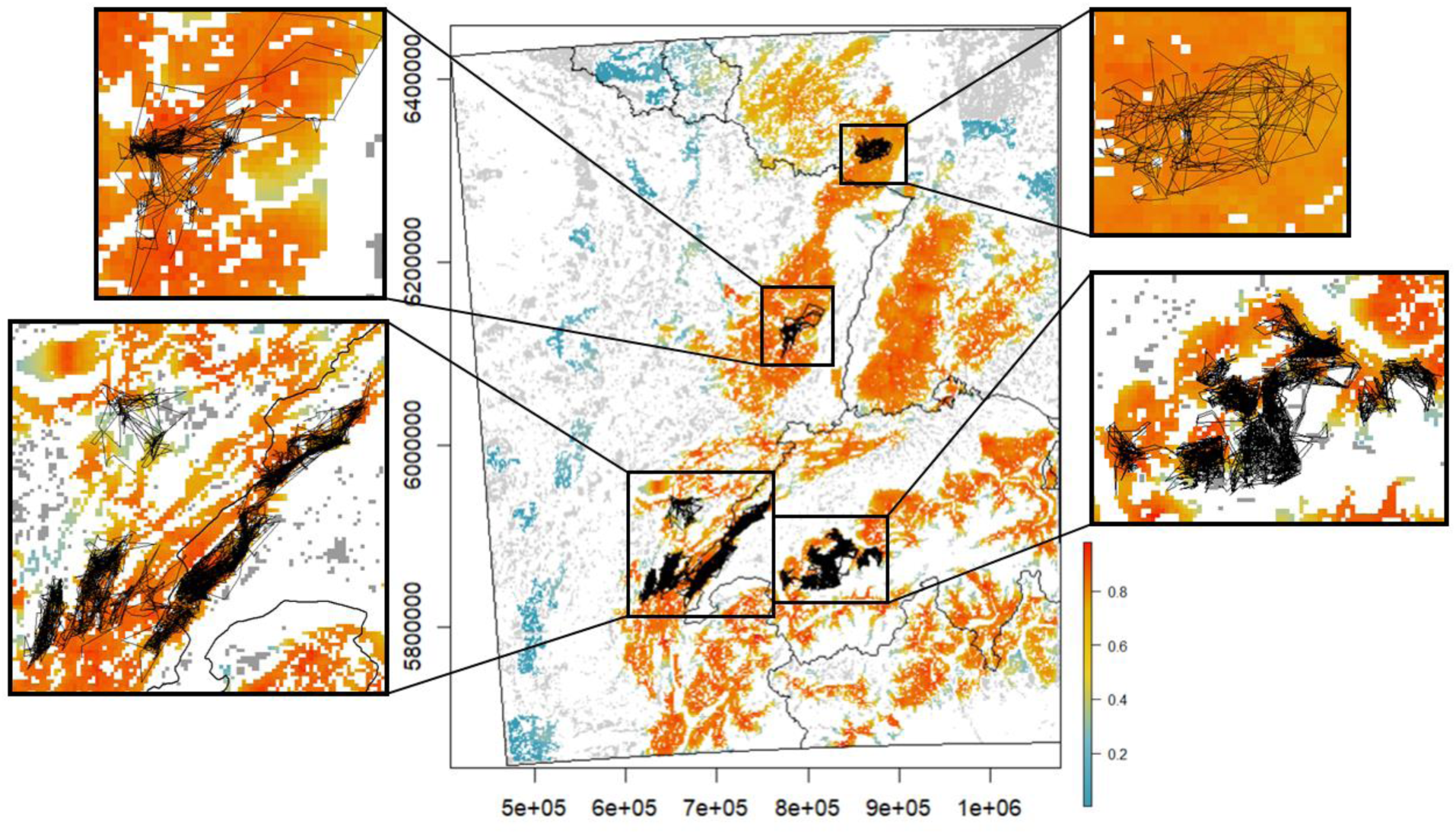
Occupancy by female territories over the study area at the last year of simulation. Values between above zero and one (color scale blue-yellow-red) are mean occupancy probability per cell of 1 km^2^ over 100 replicates (e.g., a cell equal to 1 means the cell was included in a female territory, at the 50^th^ year of simulation, in all 100 simulation replicates). Grey cells are “breeding habitats” never included in a female territory across all replicates (equal to zero). White cells are cells of a habitat type different than “breeding habitat” and therefore could never be included in a female territory. GPS and VHF recorded paths for female residents are overlaid as thin black lines. A zoom of different areas is presented on the sides of the central map.

